# The land-use dynamics of potato agrobiodiversity in the highlands of central Peru: a case study of spatial-temporal management across farming landscapes

**DOI:** 10.1101/585273

**Authors:** Alejandra Arce, Stef de Haan, Henry Juarez, Franklin Plasencia, Dharani Burra, Raul Ccanto, Severin Polreich, Maria Scurrah

## Abstract

In the high Andes, environmental and socio-economic drivers are transforming land use and presumably affecting the *in situ* conservation of potato (*Solanum* spp.). To monitor the use and conservation of intraspecific diversity, systematic and comparative studies across land-use systems are needed. We investigated the spatial-temporal dynamics of potato in two contrasting landscapes of Peru’s central Andes: a highland plateau (Huancavelica) vs. an eastern slope (Pasco). We examined household-level areal allocations, altitudinal distribution, sectoral fallowing practices, and the conservation status for three main cultivar groups: (i) bred varieties, (ii) floury landraces, and (iii) bitter landraces. Mixed methods were used to survey 323 households and the 1,101 potato fields they managed in 2012–2013. We compared the contemporary altitudinal distribution of landraces with 1975–1985 altimeter genebank data from the International Potato Center. We show that intensification occurs in each landscape through adaptations of traditional management practices while maintaining high intraspecific diversity. Access to land and production end use (sale vs. consumption) significantly affected smallholder management and differentiated the landscapes. Total areas in Huancavelica and Pasco were allocated to 82.9% vs. 74.2% floury landraces, 9.2% vs. 25.7% bred varieties, and 7.9% vs. 0.1% bitter landraces. In market-oriented Pasco, fields in sectoral fallows between 3,901 m and 4,116 m above sea level consistently contained the highest levels of landrace diversity. The bulk of diversity in subsistence-oriented Huancavelica occurred between 3,909 m and 4,324 m outside sectoral fallows. Most of the unique landraces documented were scarce across households: 45.4% and 61.7% respectively in Huancavelica and Pasco. Bred varieties showed the widest (1,100 m) and bitter landraces the narrowest (400 m) altitudinal distributions. Potato cultivation has moved upward by an average of 306 m since 1975. Landrace diversity is versatile but unevenly distributed across landscapes. This requires adaptive ways to incentivize *in situ* conservation.

## 1. Introduction

In the Andes, demographic shifts, migration, part-time farming, market integration, urbanization and climate change will increasingly affect the land-use systems that support farmers’ on-farm agrobiodiversity and the *in situ* conservation of major food plants [1–7]. Land-use responses in the Andes to the above-mentioned drivers have been varied. In some farming environments, the intensity of land use has increased in terms of cropping frequencies and areal coverage of cash crops or bred varieties, fertilizers and pesticides driven by agricultural specialization [8–11]. Other areas have seen a mixed trend due to migration, off-farm work, land abandonment, and a livelihood shift away from subsistence agriculture [12–15]. At high altitude, the expansion of agriculture resulting from climate change and market incentives is seen to encroach upon natural habitats, disrupting ecosystem services such as the provision of soil organic carbon stocks and water, and competing with other smallholder livelihood activities [16–18]. The net outcome of these processes on farmers’ management practices involving agrobiodiversity –particularly crop landrace diversity– has not been necessarily negative, as smallholder farming systems have been shown to be highly adaptive and opportunistic [19–21]. Therefore, Andean smallholder farming systems are still recognized to harbor high levels of agrobiodiversity essential for adaptive agriculture and food security [22–24].

Modern-day environmental, demographic, and socio-economic changes are nonetheless demanding ever more complex land-use choices from smallholder farmers. Processes of intensification reflect hybrid systems where traditional management schemes coexist with management modifications [25–28]. Contemporary agricultural land-use change in the high Andes is often associated with an upward expansion of cropping, micro-fragmentation of household cropping areas, incremental occurrence of pests and disease at higher altitudes, and the gradual abandonment of communal land-use management such as sectoral fallowing systems [6, 29–31]. Mixed livestock–crop systems, and competition between these two components, are particularly common at high altitudes [17, 32]. Nonetheless, it is difficult to make generalizations about many of these processes in the region due to its socioeconomic and agroecological diversity [33, 34]. The co-existence of traditional and modern management practices is not uncommon as smallholders adjust their livelihoods by integrating into markets and adopting new technologies [10, 19, 35, 36].

The persistence of high crop and landrace diversity in the portfolios of smallholder farmers has been considered a unique feature of Andean agriculture despite accelerated change, although in-depth inquiries into the relationship of land-use change and intraspecific diversity of crops are scant. In the central Peruvian highlands, potato agriculture has evolved in a harsh and risk-prone mountain environment. Its diverse microclimates, altitudinal gradients and soil conditions have led to spatially heterogeneous farming landscapes and a suite of management adaptations involving different tillage systems and field scattering, among other practices [37–39]. Extreme and typically localized weather events like frost and hail regularly result in crop failure [40]. Pest and disease outbreaks are also known to occasionally affect these high-altitude farming environments [41, 42]. To mitigate imminent risk and safeguard their food reserves and seed stocks, farmers have developed practices that juxtapose spatial and temporal features of land use at household and communal levels.

An example involves the sectoral fallowing system, or *laymi* in Quechua, as it aggregates households’ individually assigned fields into six to 10 sectors and is collectively cultivated following a crop–pasture rotation regimen [43–45]. Sectoral fallowing systems allow fragile high-altitude soils to partially recover their fertility while making pastureland available for grazing animals [46]. They also optimize labor through community-level coordination [47, 48]. Yet another example involves distinct types of tillage systems for potato cultivation [38]. *Chiwa* is a low-labor-intensity minimal-tillage practice and is commonly applied in sloping environments reserved for landraces. *Chacmeo* is another minimum-tillage practice that is moderately labor-intensive and well adapted to slope planting of landraces. *Barbecho* is a full-tillage practice and labor-intensive. It is commonly used for market-oriented production of bred varieties and commercial landraces.

Adaptive land-use practices have thus enabled smallholder farmers in Peru’s central Andes to manage high intraspecific diversity of the potato. Four botanical species of cultivated potato are recognized following the latest taxonomic treatment: *Solanum tuberosum*, *Solanum curtilobum*, *Solanum ajanhuiri*, and *Solanum juzepczukii* [49, 50]. At the intraspecific level farmers maintain an ample repertoire of genetically and morphologically distinct, farmer-recognized landraces. These landraces –each with a farmer-recognized vernacular name– are the basic unit of management and conservation on the farm [51, 52]. At the national level this intraspecific diversity is high and consists of an estimated 2,800 to 3,300 potato landraces [53]. Even at the village and household levels, landrace diversity can be remarkable. For example, in one hotspot of potato diversity, up to 406 genetically distinct landraces have been identified in the landrace portfolios of just eight farmer households, and individual households are known to maintain as many as 160 unique landraces [54].

Farmers predominantly classify cultivar groups, varieties or landraces according to visual phenotypic characters [55, 56]. Three main cultivar groups are recognized by smallholder farmers in Peru’s central highlands. The floury landraces (*S. tuberosum* Andigenum Group), also known as “boiling potatoes”, are deemed of high culinary quality and make up the bulk of the potato landrace diversity managed by farmers. They are most often cultivated as mixed lots (*chalo*, *chaqru* or *waychuy* in Quechua) containing between four and 80 floury landraces while a minority (i.e. eight landraces) are commercially produced in single-cultivar fields [57]. Bitter landraces (*S. juzepczukii* and *S. curtilobum*) are generally frost-resistant and only apt to be consumed as freeze-dried *chuño* due to their high glycoalkaloid content [40, 58]. They are also less diverse in number compared to floury landraces. Bred varieties (*S. tuberosum*) are the result of formal breeding programs and have been amply disseminated for their high-yield and disease-resistance traits in Peru. Farmers have widely integrated these into their cropping portfolios. Bred varieties occupy a special window in terms of food supply as they produce earlier than the floury landraces. They serve a dual purpose: consumption and the market.

Research concerning the contemporary spatial management of Andean smallholders’ agrobiodiversity, and specifically the interaction between land use and intraspecific diversity, can help to gain insights into multilevel conservation within and among landscapes, households and fields. In this in-depth case study, we scrutinize the land-use dynamics of the potato in two distinct diversity hotspots in Peru’s central Andes. We examine and compare areal allocations, altitudinal ranges, fallowing rates, the use of sectoral fallowing, and the conservation status of individual landraces. To detect possible temporal changes in the distribution of landraces, we compare the contemporary altitudinal range with 1975–1985 elevation records of accessions from the International Potato Center (CIP). We hypothesize that the spatial-temporal dynamics characterizing each landscape in the central Peruvian highlands is driven by context-specific pressures that require smallholders’ differential management adjustments while allowing the maintenance of high intraspecific diversity. Implications for the long-term *in situ* conservation tied to land use are reflected upon.

## 2. Materials and methods

### 2.1. Study area and household sample

We conducted in-depth research in five communities pertaining to two contrasting highland landscapes of Peru’s central Andes (Fig 1; Table 1). The first cluster of three farmer communities lies in the central plateau or cordillera of the Huancavelica region where potato is grown at high altitude with frequent exposure to frost and hail. The second cluster of two communities is nestled in a valley along the eastern flanks of the Andes in the Pasco region, about 235 kilometers from the Huancavelica region. Here relatively humid conditions lead to high levels of pressure from late blight disease (*Phytophthora infestans*). Farmers in Huancavelica are indigenous Quechua speakers, while those in Pasco are mostly mestizo Spanish speakers. Both sites are recognized hotspots of potato intraspecific diversity [59, 60]. A total of 176 and 147 households in the Huancavelica and Pasco landscapes, respectively, were randomly sampled and participated in the study.

**Fig 1.**
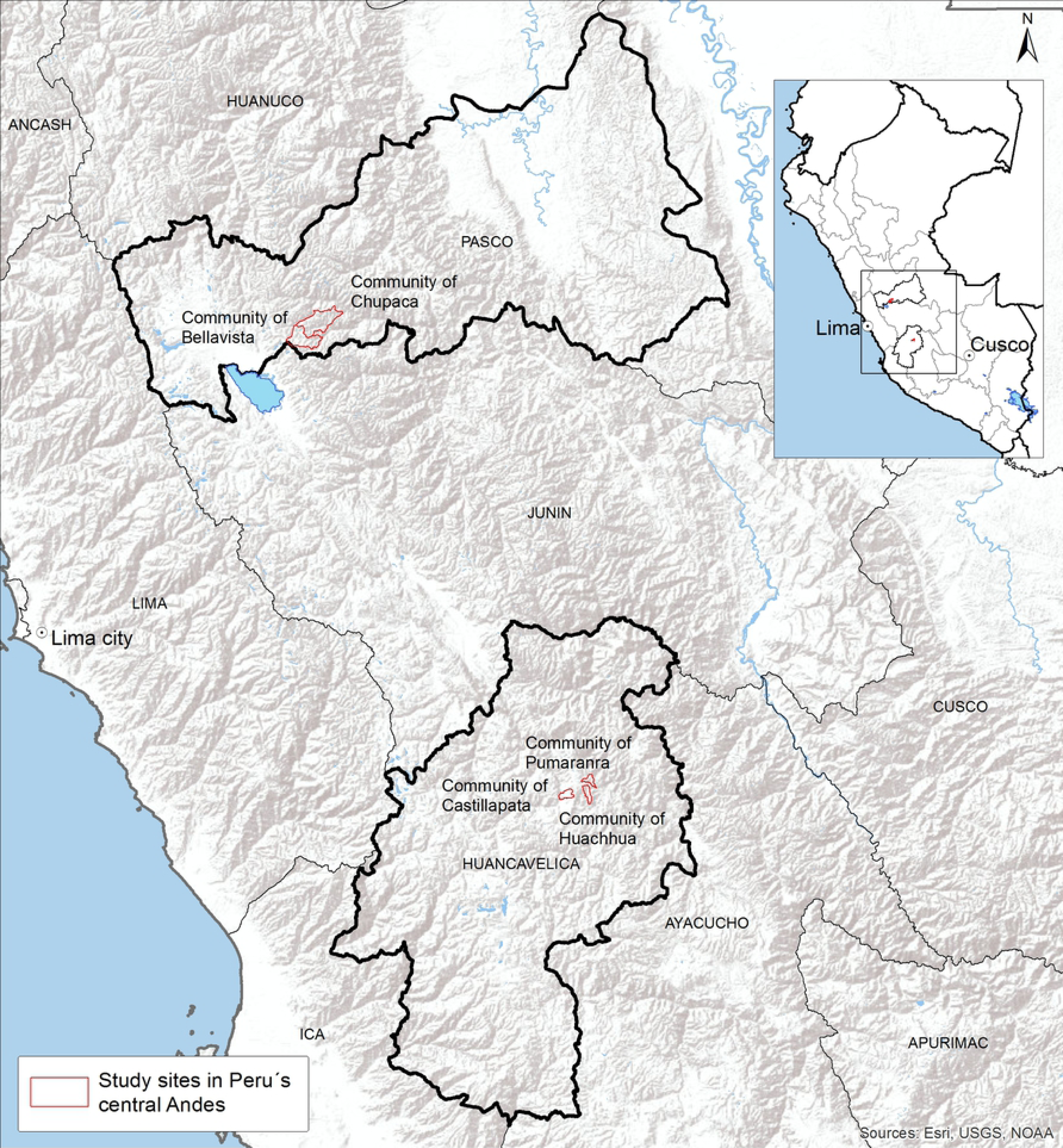
Study sites in Peru’s central Andes.

**Table 1.**
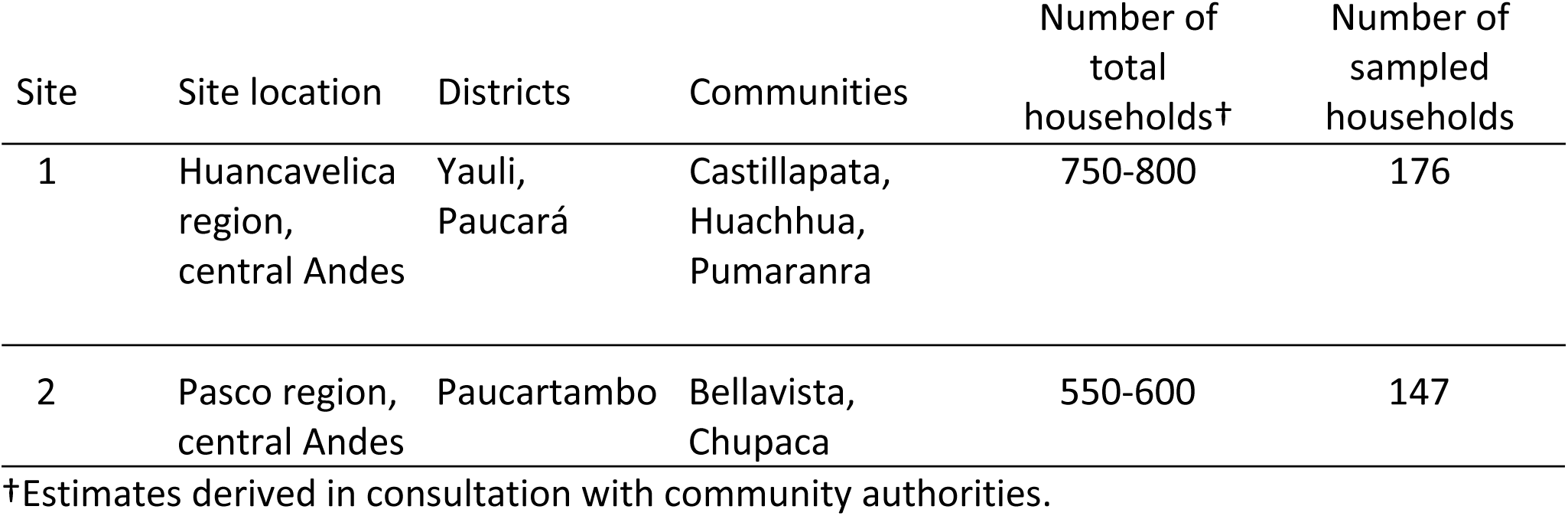
Study sites in Peru’s central Andes.

### 2.2. Participatory mapping and field-level sampling

Drawing from cartography and participatory methods we conducted participatory mapping (pGIS) between February and June 2013 to document the land use of each potato field of participating households. The procedure consisted of two parts. First, we accompanied farmers on one or two visits to each of their potato fields for short surveys, georeferencing, and field sampling of cultivars planted. Second, we ran multiple focus-group meetings centered on drawing over printed high-resolution satellite images of each of the five communities. Participating households located and drew each of their potato fields on the base map. Local authorities delimited community boundaries and identified each of the sectors comprising fallowing systems.

Field-level surveys were conducted with each household (n=323). Trained enumerators implemented the surveys in Quechua (Huancavelica) and Spanish (Pasco). Each survey had four components: (i) basic household-level information, (ii) field-level characteristics of each potato field, (iii) georeferencing each potato field with Garmin Oregon 550t global positioning systems (GPS) devices, and (iv) cultivar diversity sampling at harvest. For each georeferenced field a range of variables was collected, including planting date, fallowing-sector association, tillage type, use of chemicals, slope, seed source, and product end use. Georeferencing resulted in the collection of waypoints for the corners and center of each field, as well as altitude. Farmers also recalled crop species content and fallows for each year from 2004 to 2013. A total of 1,101 potato fields, 481 in Huancavelica and 620 in Pasco, were visited, surveyed and georeferenced.

During the potato harvest from April to June 2013, each potato field (n=1,101) was sampled for its cultivars. In each field, we randomly selected 25 potato plants that were distributed along eight equidistant rows and unearthed one tuber per plant until we arrived at a total count of 200 tubers. In cases where the household had already harvested, we randomly picked 200 tubers from the heap or bags. The sampled tubers served to identify and count each of the individual cultivars following the local nomenclature used by farmers. This exercise was carried out by local survey teams and the farmers to whom each field belonged. In each field, the occurrence of a potato cultivar was recorded as the total count of individual tubers out of 200 total tubers sampled.

### 2.3. Focus-group meetings to refine cultivar classification

Individual cultivars are frequently recognized by more than one name (synonyms), and sometimes the same name is used for distinct cultivars (homonyms). This poses a challenge of over- or under-classification [51]. To overcome this issue, we carried out focus group meetings with farmers who were the most knowledgeable about varietal diversity. A representative collection of the distinct cultivar morphotypes that were identified during field surveys was created for each community by using real tuber samples and, in a few cases, photographs. Local experts, both men and women, indicated alternate names associated with each tuber sample. A list of unique cultivars and their synonyms was thus derived for each community. These, in turn, were compared and cross-checked for the same tuber samples for each landscape. A master list of unique cultivars was attained for each of the two landscapes.

### 2.4. Conservation status of cultivars

To determine the conservation status of cultivars for each landscape (Huancavelica, Pasco) we used two indices (59): (i) relative cultivar frequency (RCF), (ii) overall cultivar frequency (OCF). The RCF index is used to gauge the relative abundance or frequency (or rarity) of a unique cultivar in comparison to all other cultivars sampled in each landscape It indicates the proportion of each distinct cultivar over the total cultivar population sampled in each landscape. For each cultivar occurrence per household, a household cultivar frequency (HCF) was first calculated. This involved summing the number of tubers sampled for a specific cultivar across a household’s total fields, dividing the result by the total number of samples of all cultivars for that household, and multiplying by 100%. The RCF for each cultivar was then derived by summing its corresponding HCFs and dividing the result by the total number of households sampled per landscape. Red listing was based on the threshold levels: RCF<0.05=very scarce, RCF<0.10=scarce, RCF<0.25=uncommon, RCF<1.00=common, RCF>1.00=abundant.

The OCF index is a measure of evenness. For each cultivar, its community cultivar frequency (CCF) was first calculated by dividing the number of households cultivating it by the total number of sampled households in each community comprising a landscape and multiplying by 100%. The OCF for each cultivar was obtained by summing its CCFs and dividing the result by the total number of communities sampled in the landscape. The evenness of individual cultivars was then classified as the proportion of households growing them: OCF<1%=very few households, OCF<5%=few households, OCF<25%=many households, OCF>25%=most households.

### 2.5. Timeline series analysis

Possible changes in the altitudinal distributions of floury and bitter landraces were examined. We compared the altitudes documented in this study with genebank passport altimeter data from all collections made in 1975–1985 for the same two landscapes. The latter data were provided by the International Potato Center and totaled 63 georeferenced landrace accessions from 16 locations in Huancavelica and Pasco.

### 2.6. Statistical analyses

Descriptive statistical analyses were performed using the statistical computing software R version 3.4.1 [61]. Household averages for number of potato cultivars, number of fields, and cropping areas were calculated by cultivar group (bred varieties, floury landraces, bitter landraces) and landscape. For each landscape, we calculated the potato cropping area by cultivar group and altitudinal distribution range in total number of hectares. We examined the number of fields, the number of cultivars per field, and fallowing rates per field by cultivar group for each landscape. Fallowing rates were obtained by dividing the number of unplowed (fallow) years by the total number of years included in the cropping cycle. We analyzed changes in the altitudinal distribution of floury and bitter landraces from 1975 to 2013 by calculating their average, maximum, minimum, and standard deviation values. To detect significant differences in the number of cultivars, areas, and altitudes between fields associated with a fallowing sector and not associated with a sector within landscapes we performed two-sample unpaired Wilcoxon tests. Significance was determined at the p<0.001 level. To identify salient distinctions in farmers’ field management practices between landscapes, fields were classified as low, intermediate, or high-range, based on their altitude. The altitudinal range for low was 3,097-3,499 m, for intermediate 3,500-3,899 m, and for high 3,900-4,324 m. This classification resulted in 97 intermediate-range and 382 high-range fields in Huancavelica, and 379 intermediate-range and 207 high-range fields in Pasco. For each high and intermediate range, several regression and statistical learning approaches were compared, and the best-performing model (details below) was used to identify management characteristics that significantly differentiated fields across landscapes. This analysis was not carried out for low-range fields as they were too few (two in Huancavelica and 34 in Pasco) to compare between landscapes. Models using logistic regression, generalized linear models (using lasso, elastic and ridge-based penalized maximum likelihood approaches) and random forest-based approaches were built using field-level management practices data (i.e. cultivar group content, number of cultivars, field area, days to harvest, planting season, sector association, seed source, product end use, tillage type, application (yes/no) of chemicals, and fallowing rate) collected for each field surveyed as explanatory variables, and landscapes as the outcome variable.

Receiver operating characteristic (ROC), sensitivity and specificity metrics with ten-fold cross validation were used to assess model quality. The coefficient of variation metric was used to identify the lowest lambda value for lasso and ridge-based penalized general linear models. To account for imbalance in the number of intermediate-range fields (97 in Huancavelica and 379 in Pasco), up and down sampling approaches were employed to build the models. The generalized linear model with elastic-based penalization approach was found to perform best in classifying intermediate-range fields and the generalized linear model with ridge-based penalization approach performed best in classifying high-range fields across landscapes. The above analysis was performed in the R statistical computing environment using the packages glmnet caret and catools [62]. The outputs of the models were visualized through boxplots drawn with the ggplot2 package, and association plots (based on an independence model and Pearson test of the residuals) were drawn using the vcd package in the R statistical computing environment [61, 63, 64].

Logistic regression was performed to identify significantly different household-level characteristics between landscapes. The variables age and sex of the household head, number of children and adults in the household, total number of potato fields for the household, off-farm income (yes/no), total number of bred varieties, floury landraces and bitter landraces across all fields belonging to the household, and average household area under bred, floury, and bitter cultivation were used as explanatory variables, with landscapes serving as the outcome variable. Stepwise regression (forward and backward) was employed, and the resulting model was selected based on Akaike information criterion (AIC) and likelihood ratio test (LRT) criteria. Statistical analysis of field-level cropping history and land-use patterns (2004–2013) was performed using R package TraMineR to elucidate differences between landscapes at each altitudinal range (intermediate and high) separately [65].

### 2.7. Research ethics

The study was conducted in accordance with the Declaration of Helsinki and the guidelines provided by the Central Committee on Research Ethics at the University of Antioquia, Medellín. Ethics approval was not required for this research according to national regulations as it involved human subjects in non-invasive survey procedures. We sought and obtained the approval of community authorities prior to survey implementation. We described the objectives of the study, the methodology, the oral prior informed consent option, voluntary nature and confidentiality of households participating during a community assembly. Community authorities from the five communities selected agreed to the study. Households were surveyed only after community-level approval.

## 3. Results

### 3.1. Household characteristics

We calculated and compared main household features across landscapes (Table 2). These indicated demographic and socio-economic distinctions, such as in the average number of children per household, the proportion of heads of household without formal schooling, and family vs. hired labor to sustain agricultural activities on the farm. The most significant differences between households in Huancavelica and Pasco as detected by logistic regression analyses (best model) were number of children, number of fields, off-farm income, number of floury landraces, and average area cultivated with bred varieties (Table 3).

**Table 2.** Main household-level characteristics by landscape.

| **Demography** | **Huancavelica (n†=176)** | **Pasco (n†=147)** |
| --- | --- | --- |
| Average age of head of household (years) | 47.7 (±15.0) | 44.3 (±14.3) |
| Female heads of household (%) | 10.0 | 8.0 |
| Average number of children (<18 years) per household | 2.4 (±2.0) | 1.4 (±1.2) |
| Average number of total household members per household | 4.7 (±2.3) | 3.8 (±1.5) |
| **Education** |  |  |
| Heads of household who completed primary education (%) | 8.0 | 23.1 |
| Heads of household who did not complete primary education (%) | 31.2 | 31.9 |
| Heads of household who completed secondary education (%) | 19.9 | 13.6 |
| Heads of household who did not complete secondary education | 12.5 | 25.2 |
| Heads of household who attended technical school or college (%) | 4.0 | 1.4 |
| Heads of household who did not have any formal schooling (%) | 24.4 | 4.8 |
| **Sources of farm labor** |  |  |
| Family only (%) | 46.6 | 23.1 |
| Family and reciprocity (%) | 35.8 | 23.8 |
| Family and hired labor (%) | 3.4 | 35.4 |
| Reciprocity and communal work (%) | 6.3 | 9.5 |
| Family, hired and reciprocity (%) | 3.4 | 2.7 |
| Hired labor (%) | 2.8 | 4.1 |
| Hired and reciprocity or communal work (%) | 1.7 | 1.4 |
| **Potato cropping** |  |  |
| Households planting bred varieties (%) | 64.8 | 78.9 |
| Households planting floury landraces (%) | 99.4 | 100.0 |
| Households planting bitter landraces (%) | 39.8 | 3.8 |
| **Off-farm income** |  |  |
| Households with off-farm sources of income (%) | 60.8 | 68.7 |

†Total number of households per landscape.

**Table 3.** Logistic regression output (best model) of most significant differentiating household characteristics between the Huancavelica and Pasco landscapes.

| **Significant explanatory variables†** | **Odds ratio** | **2.50%** | **97.50%** |
| --- | --- | --- | --- |
| (Intercept) | 0.2401 | 0.0895 | 0.6169 |
| Number of children per household | 0.6490 | 0.5250 | 0.7883 |
| Number of fields per household | 1.9775 | 1.6270 | 2.4605 |
| Off-farm income | 3.4088 | 1.7822 | 6.7795 |
| Number of floury landraces | 0.9272 | 0.9001 | 0.9519 |
| Average area cultivated with bred varieties | 1.0012 | 1.0003 | 1.0023 |

†Significant explanatory variables correspond to variables used in the logistic regression model that were identified to significantly differentiate households in Pasco from those in Huancavelica. The odds ratio was calculated by exponentiating the coefficients (of significant variables) obtained from the logistic regression model, while the columns 2.5% and 97.5% correspond to the exponentiated confidence interval levels.

### 3.2. Field-management characteristics

The number of potato fields cropped per household was 2.7 (±1.4) in Huancavelica and 4.3 (±2.1) in Pasco. Rented fields represented 11.9% of total fields only in Pasco. Potato production in Huancavelica was destined for household consumption for 78.0% and dual purpose (consumption and sale) for 22.0% of fields. In Pasco, production for sale represented 60.0%, dual purpose 23.5%, and solely consumption 16.5%. Most field production had a secondary end use. In Huancavelica, farmers saved medium-sized tubers for both seed and making freeze-dried *chuño* from 90.7% of fields. Seed and *chuño* production exclusively were secondary uses for 8.1% and 0.4% of fields respectively. Only 0.8% of production from sampled fields had no secondary end use. In Pasco, secondary uses were seed and *chuño* production (20.0%), tuber seed exclusively (39.4%), *chuño* production exclusively (28.4%), seed and pig feed (4.8%), pig feed exclusively (1.1%), *chuño* and pig feed (0.8%). Only 5.5% of production from surveyed fields did not have any secondary end use.

In both landscapes, households followed two potato cropping calendars, the *qatun tarpuy*, literally ‘big planting’ (main season), and the *michka*, or small planting (off-season). The ‘big plantings’ coincide with the main rainy season and span from October-November (sowing period) to May-June (harvesting period). It is the most intensive season in terms of labor demands. The off-season plantings are short, involve small cropping areas and generally demand access to irrigation with sowing taking place from June to July (dry season). Consequently, most potato fields mapped corresponded to the main season: 97.1% and 82.4% of fields in Huancavelica and Pasco respectively. The number of main and off-season fields per household, respectively, was 2.7 (±1.3) vs. 0.1 (±0.2) in Huancavelica, and 3.5 (±1.9) vs. 0.8 (±0.9) in Pasco. Pasco had the longer potato-growing calendar. The number of days to harvest was 261.9 (±32.1) compared to 197.3 (±21.7) in Huancavelica. However, the minimum and maximum number of days to harvest recorded for each were similar: 121 and 304 in Huancavelica vs. 120 and 309 in Pasco, depending on the cultivar group and specific cultivar involved.

All potato fields in Pasco and 44.7% of fields in Huancavelica received applications of chemicals (fungicides and fertilizers). Most potato fields, 71.9% in Huancavelica and 100% in Pasco, were managed with the *chiwa* tillage system, followed by *barbecho* (22.5%) and *chacmeo* (5.6%) in Huancavelica. In this central plateau, fields with floury landraces were tilled 73.1% *chiwa*, 23.2% *barbecho* and 3.7% *chacmeo*; fields with bred varieties were tilled 68.8% *chiwa*, 22.4% *barbecho* and 8.8% *chacmeo*; and fields with bitter landraces were tilled 95.2% *chiwa*, 1.9% *barbecho* and 2.9% *chacmeo*.

### 3.3. Cultivar diversity, abundance and evenness

Field sampling and focus group meetings resulted in the identification of 130 and 191 unique cultivars for Huancavelica and Pasco respectively. Floury landraces represented the bulk of diversity: 85.5% of cultivars in Huancavelica and 95.8% in Pasco. Bred varieties made up 9.2% and bitter landraces 5.3% of cultivars in Huancavelica. In Pasco, bred varieties were 3.7% and bitter landraces 0.5% of cultivar diversity. Floury landraces dominated households’ portfolios (Table 4). The maximum number of cultivars for any household (56) was recorded for this cultivar group in Pasco. Bred and bitter landraces registered a maximum household-level cultivar count of 6 and 5 cultivars respectively in Huancavelica.

**Table 4.** Number of distinct cultivars managed per household by cultivar group and landscape.

| Huancavelica |  |  |  |  |  | Pasco |  |  |  |  |  |
| --- | --- | --- | --- | --- | --- | --- | --- | --- | --- | --- | --- |
| Cultivar group | N† | Av. | Max. | Min. | SD | Cultivar group | N† | Av. | Max. | Min. | SD |
| Bred varieties | 114 | 1.8 | 6.0 | 1.0 | 1.1 | Bred varieties | 116 | 1.4 | 4.0 | 1.0 | 0.7 |
| Floury landraces | 175 | 12.5 | 42.0 | 1.0 | 7.1 | Floury landraces | 147 | 16.5 | 56.0 | 1.0 | 11.0 |
| Bitter landraces | 70 | 1.7 | 5.0 | 1.0 | 1.1 | Bitter landraces | 5 | 1.0 | 1.0 | 1.0 | 0.0 |
| Total | 176 | 14.3‡ | 49‡ | 1‡ | 8.0‡ | Total | 147 | 17.7‡ | 58‡ | 2‡ | 11.1‡ |

†Number of households planting each cultivar group.
‡Calculated from sum of distinct cultivars across the three cultivar groups.

We contrasted the spatial distribution and relative abundance of cultivars by cultivar group (Fig 2A, 2B) and RCF level (Fig 3A, 3B) for a representative community in each landscape. Red listing showed that most cultivars were very scarce (RCF<0.05) across households: 45.4% of total cultivars in Huancavelica and 61.7% in Pasco (Table 5). These were predominantly floury landraces. Huancavelica showed comparatively more common and abundant cultivars than Pasco. In terms of evenness, approximately two thirds of cultivars in each landscape were grown by very few households (OCF<1%) or few households (OCF<5%) while less than 15% of cultivars were present in the cropping portfolios of most households (OCF>25%; Table 6). Overall, for the landscapes combined, 12.5% of cultivars were in the cropping portfolios of most households while 29.6% were grown by less than 1% of households.

**Fig 2A.**
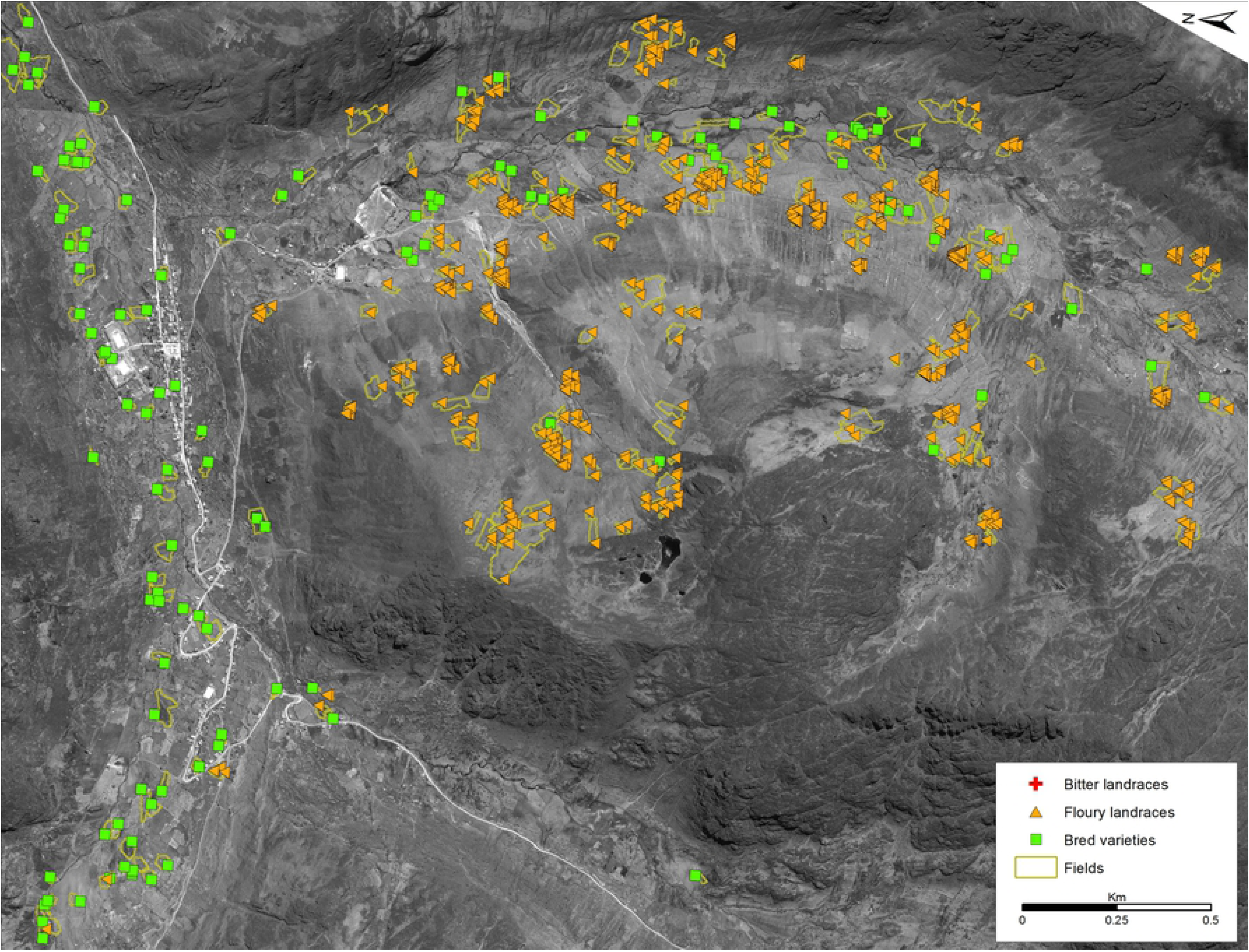
Spatial distribution of bitter landraces, floury landraces and bred varieties in the community of Bellavista, Pasco.

**Fig 2B.**
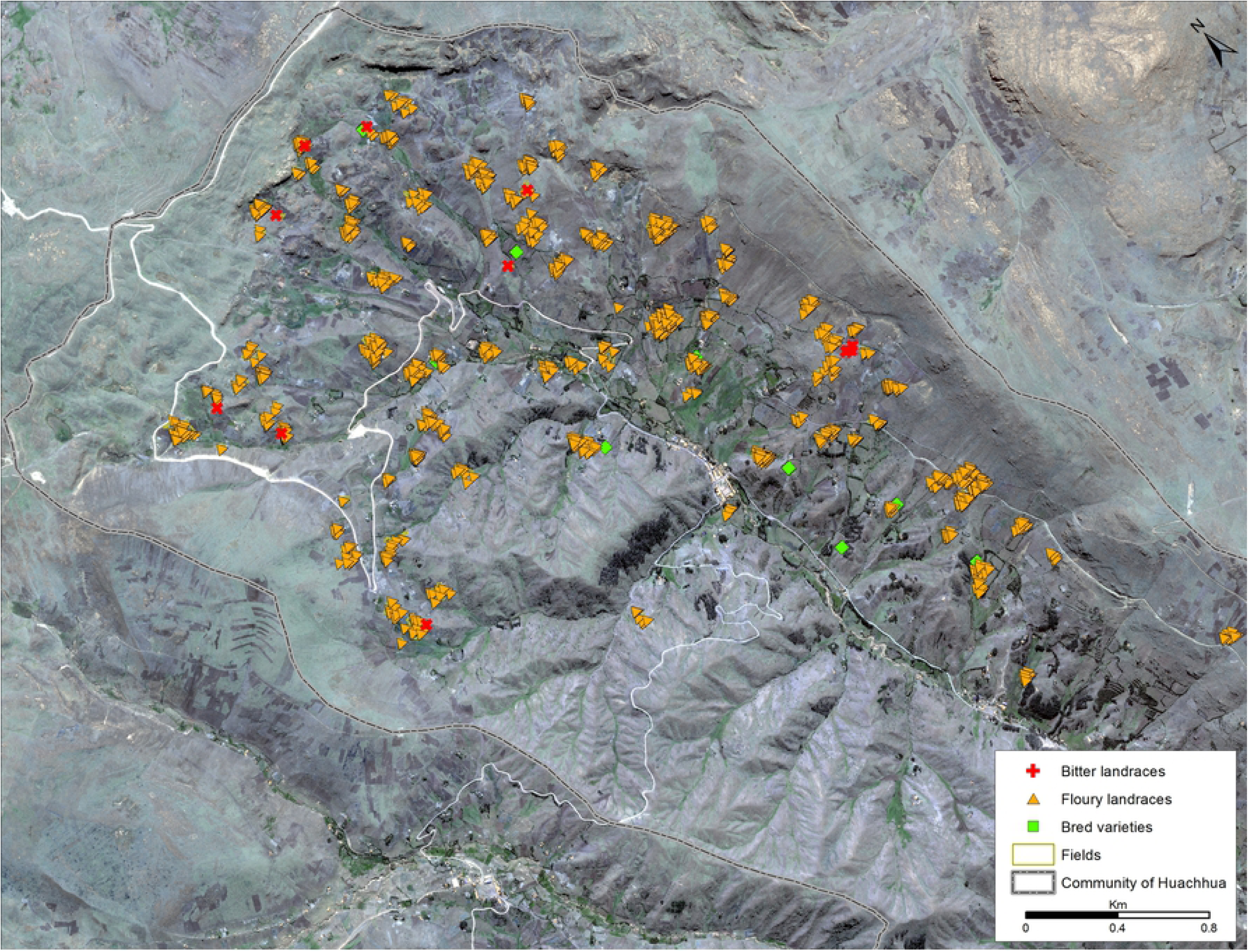
Spatial distribution of bitter landraces, floury landraces and bred varieties in the community of Huachhua, Huancavelica.

**Fig 3A.**
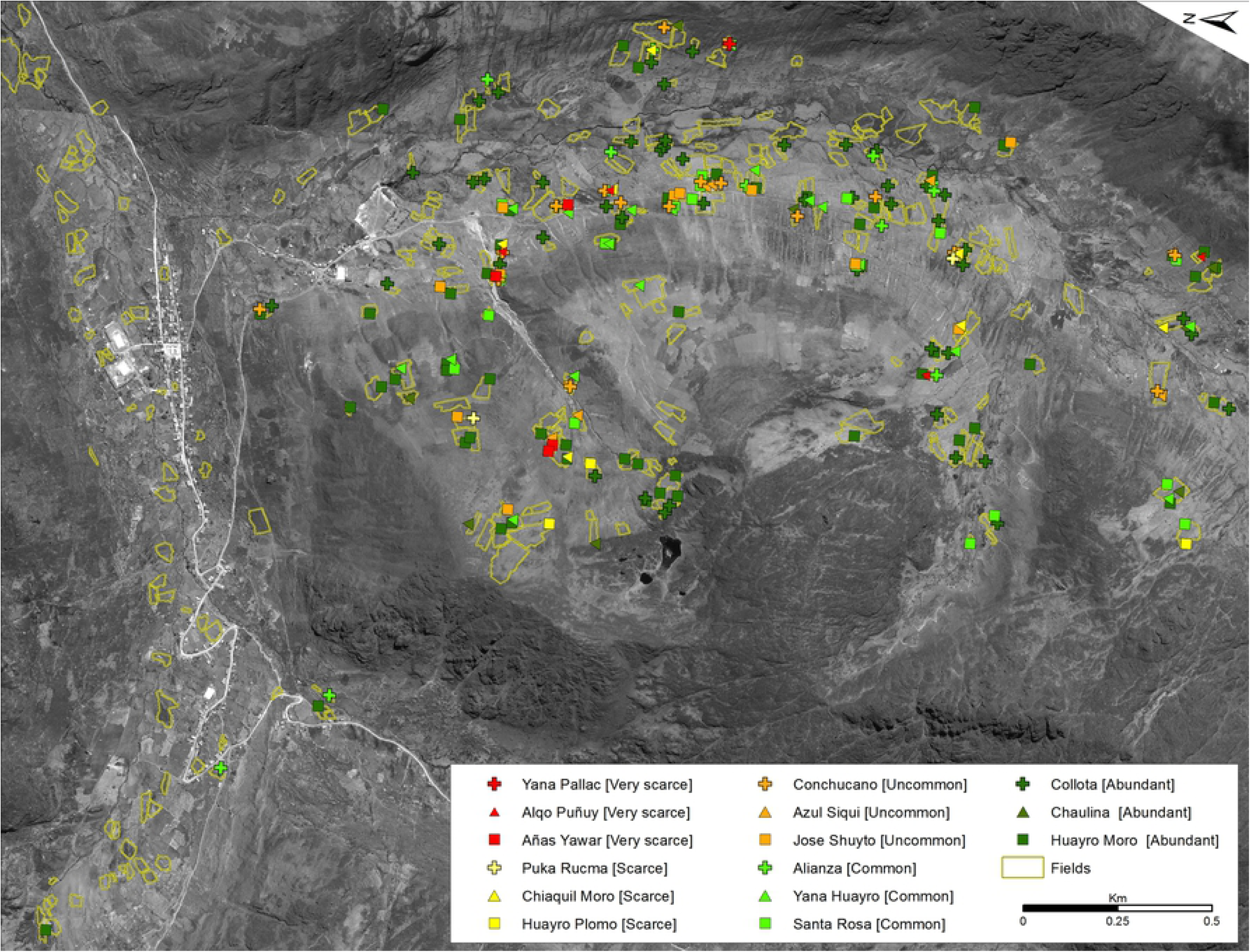
Spatial distribution of cultivars selected for their top RCF index values in each RCF level (very scarce, scarce, uncommon, common, abundant) in the community of Bellavista, Pasco.

**Fig 3B.**
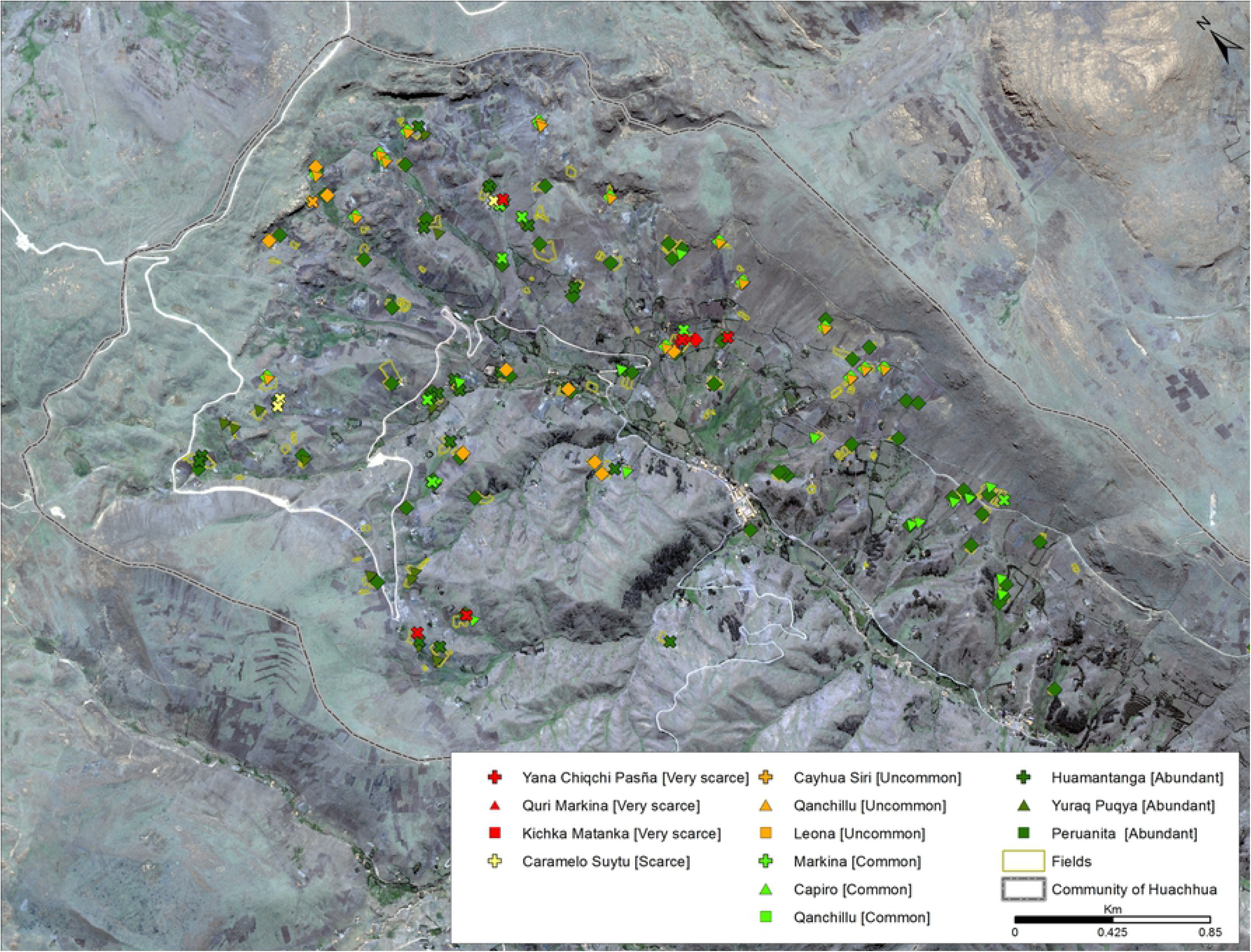
Spatial distribution of cultivars selected for their top RCF index values in each RCF level (very scarce, scarce, uncommon, common, abundant) in the community of Huachhua, Huancavelica.

**Table 5.** Relative cultivar frequencies (RCF) or measure of relative abundance of cultivars by cultivar group and landscape.

| Huancavelica |  |  |  |  |  |  |  |  |  |  |
| --- | --- | --- | --- | --- | --- | --- | --- | --- | --- | --- |
|  | Very scarce (<0.05) |  | Scarce (<0.10) |  | Uncommon (<0.25) |  | Common (<1.00) |  | Abundant (>1.00) |  |
| Cultivar group | No. of cultivars | %* | No. of cultivars | % | No. of cultivars | % | No. of cultivars | % | No. of cultivars | % |
| Bred varieties | 3 | 2.3 | 0 | 0.0 | 1 | 0.8 | 5 | 3.8 | 3 | 2.3 |
| Floury landraces | 55 | 42.3 | 8 | 6.2 | 11 | 8.5 | 22 | 16.9 | 15 | 11.5 |
| Bitter landraces | 1 | 0.8 | 0 | 0.0 | 1 | 0.8 | 2 | 1.5 | 3 | 2.3 |
| Total† | 59 | 45.4 | 8 | 6.2 | 13 | 10.1 | 29 | 22.2 | 21 | 16.1 |

| Pasco |  |  |  |  |  |  |  |  |  |  |
| --- | --- | --- | --- | --- | --- | --- | --- | --- | --- | --- |
|  | Very scarce (<0.05) |  | Scarce (<0.10) |  | Uncommon (<0.25) |  | Common (<1.00) |  | Abundant (>1.00) |  |
| Cultivar group | No. of cultivars | % | No. of cultivars | % | No. of cultivars | % | No. of cultivars | % | No. of cultivars | % |
| Bred varieties | 2 | 1.0 | 0 | 0.0 | 2 | 1.1 | 0 | 0.0 | 3 | 1.6 |
| Floury landraces | 116 | 60.7 | 20 | 10.5 | 22 | 11.5 | 15 | 7.9 | 10 | 5.2 |
| Bitter landraces | 0 | 0.0 | 0 | 0.0 | 1 | 0.5 | 0 | 0.0 | 0 | 0.0 |
| Total† | 118 | 61.7 | 20 | 10.5 | 25 | 13.1 | 15 | 7.9 | 13 | 6.8 |

\*Percent of total number of cultivars registered within each landscape: 130 in Huancavelica and 191 in Pasco.
†Total number of cultivars under each RCF category.

**Table 6.**
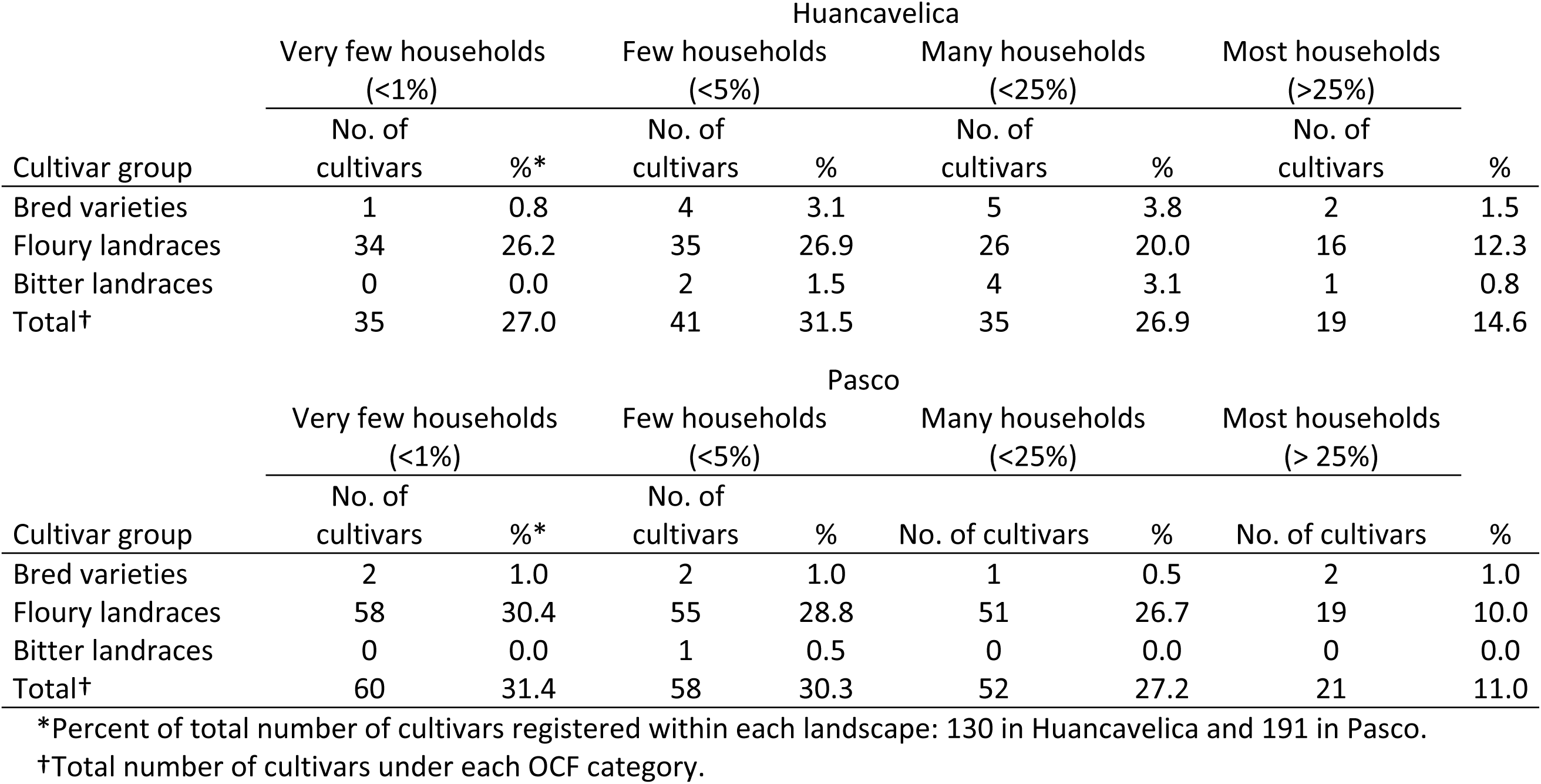
Overall cultivar frequencies (OCF) or measure of evenness of unique cultivars by cultivar group and landscape.

### 3.4. Spatial management of intraspecific diversity

#### 3.4.1. Fields with one type of cultivar compared to fields with mixed groups

Mixed fields with two to three cultivar groups contained the highest average number of distinct cultivars: 13 (±8.8) cultivars per field in Huancavelica and 14 (±6.4) in Pasco. The distribution of distinct cultivar groups within such mixed fields always involved separated sub-plots assigned to floury landraces, bitter landraces or bred varieties. Fields containing all three cultivar groups only made up 5.4% of the fields sampled in Huancavelica. In Pasco, most mixed fields comprised combinations of floury and bred cultivars and represented 11.5% of all sampled fields. These contained an average of 11.8 (±11.6) cultivars per field. Bred varieties and floury landraces occurred together in 23.1% of fields in Huancavelica, with an average of 10.2 (±5.4) cultivars per field. Across landscapes, most fields were planted exclusively with floury landraces: 48.9% of fields in Huancavelica and 60.6% in Pasco with 57.9% and 49.5% of these, respectively, containing *chaqru* mixtures of at least four cultivars. On average, exclusively floury fields contained 6.0 (±5.5) cultivars per field in Huancavelica and 6.0 (±6.8) in Pasco. A much lower proportion of fields contained exclusively bred varieties: 6.9% in Huancavelica and 27.1% in Pasco, with an average of 1.1 (±0.3) varieties per field in each landscape. Floury and bitter landraces occurred together in 11.4% of fields in Huancavelica and 0.6% in Pasco. Only in Huancavelica were fields planted exclusively with bitter landraces (4.4%) at an average 1.3 (±0.7) cultivars per field. In Pasco bitter landraces were grown with bred varieties and floury landraces in 0.8% of fields. In these cases (n=5) only one bitter landrace was cultivated out of an average of 15.8 total cultivars per field. Floury landraces were allocated the most fields per household in both landscapes (Table 7). In Pasco, the average number of fields per household with exclusively floury landraces and exclusively bred varieties surpassed that of Huancavelica by roughly one field.

**Table 7.** Average number of fields per household for exclusive and mixed fields by cultivar group and landscape.

| Cultivar group | Huancavelica |  |  |  |  | Pasco |  |  |  |  |
| --- | --- | --- | --- | --- | --- | --- | --- | --- | --- | --- |
|  | N† | Av. | Max. | Min. | SD | N† | Av. | Max. | Min. | SD |
| Bred varieties | 32 | 1.0 | 2.0 | 1.0 | 0.2 | 90 | 1.9 | 5.0 | 1.0 | 1.0 |
| Floury landraces | 126 | 1.9 | 5.0 | 1.0 | 0.9 | 138 | 2.7 | 8.0 | 1.0 | 1.7 |
| Bitter landraces | 18 | 1.2 | 3.0 | 1.0 | 0.5 | - | - | - | - | - |
| Mixed (BR+FL)‡ | 81 | 1.4 | 4.0 | 1.0 | 0.7 | 52 | 1.4 | 3.0 | 1.0 | 0.6 |
| Mixed (FL+BL)‡ | 47 | 1.2 | 2.0 | 1.0 | 0.4 | 4 | 1.0 | 1.0 | 1.0 | 0.0 |
| Mixed (BR+FL+BL)† | 26 | 1.0 | 1.0 | 1.0 | 0.0 | 1 | 1.0 | 1.0 | 1.0 | - |

†Number of households managing each field type.
‡BR=bred varieties, FL=floury landraces, BL=bitter landraces.

#### 3.4.2. Cropping areas

The total potato cropping area differed considerably between landscapes: 35.0 ha for 176 households in Huancavelica and 81.0 ha for 147 households in Pasco. Total areal proportions by cultivar group were 82.9% vs. 74.2% for floury landraces, 9.2% vs. 25.7% for bred varieties and 7.9% vs. 0.1%, for bitter landraces in Huancavelica and Pasco respectively. On average, the total household potato cropping area was 1,989 (± 1,588) m² in Huancavelica and 5,509 (± 3,994) m² in Pasco. Households in Huancavelica tend to manage much smaller areas. Floury cultivars comparatively occupied the largest areas per household (Table 8). These were 5.9 and 2.3-fold the cropping areas of bred varieties and bitter landraces, respectively, in Huancavelica, and 4.2 and 70.2-fold the cropping areas of their counterparts in Pasco. Household field sizes were notably different between the two landscapes (Table 9). These always tended to be two to three times larger for households in Pasco for fields with bred varieties and floury landraces or a mix of these two cultivar groups.

**Table 8.**
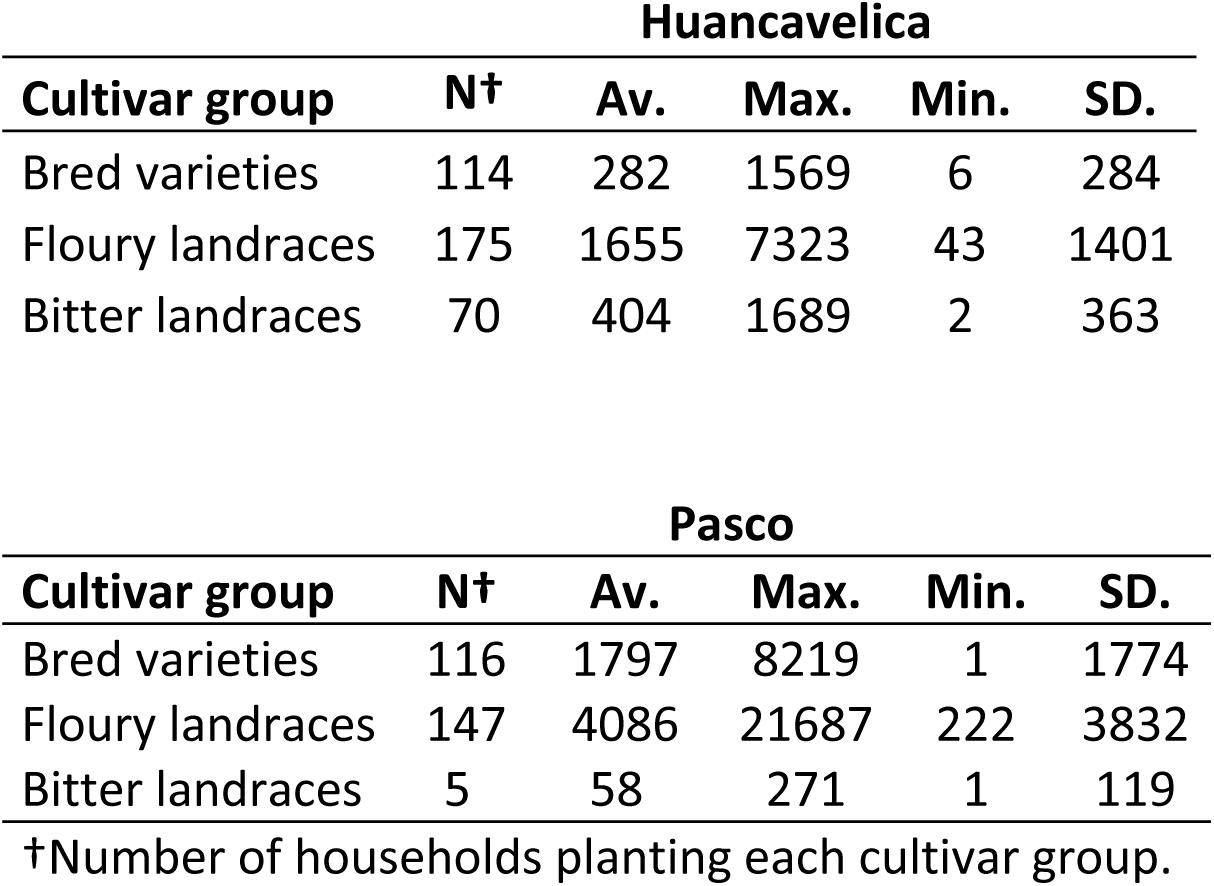
Average total cropping area (m²) per household by cultivar group and landscape.

**Table 9.** Average area (m²) per field for exclusive and mixed fields by cultivar group and landscape.

| Huancavelica |  |  |  |  |  | Pasco |  |  |  |  |
| --- | --- | --- | --- | --- | --- | --- | --- | --- | --- | --- |
| Cultivar group | N† | Av. | Max. | Min. | SD. | N† | Av. | Max. | Min. | SD. |
| Bred varieties | 33 | 340 | 1465 | 23 | 333 | 168 | 1069 | 6818 | 96 | 984 |
| Floury landraces | 235 | 627 | 3922 | 9 | 608 | 376 | 1320 | 13283 | 9 | 1562 |
| Bitter landraces | 21 | 285 | 883 | 40 | 220 | - | - | - | - | - |
| Mixed (BR+FL)‡ | 111 | 826 | 5904 | 55 | 902 | 71 | 1846 | 12917 | 44 | 2181 |
| Mixed (FL+BL)‡ | 55 | 919 | 3219 | 99 | 768 | 4 | 520 | 1375 | 17 | 613 |
| Mixed (BR+FL+BL)‡ | 26 | 1660 | 5898 | 193 | 1602 | 1 | 821 | 821 | 821 | - |

†Number of fields for each exclusive and mixed cultivar group type.
‡ BR=bred varieties, FL=floury landraces, BL=bitter landraces.

#### 3.4.3. Contemporary range of altitudes at which potatoes are grown

The altitudinal distribution of potato differed by 200 m between landscapes, with Pasco having a slightly wider range (3,000–4,200 m) and distribution in Huancavelica reaching higher altitudes (3,400–4,400 m) (Fig 4). In Huancavelica and Pasco, respectively, 84.9% and 83.5% of cultivation in terms of areal coverage occurred between 3,800 m and 4,200 m, and 3,700 m and 4,100 m. Cultivation of bred varieties and floury landraces began at 3,097 m and 3,264 m in Pasco vs. 3,464 m and 3,521 m in Huancavelica. Bred varieties and floury landraces overlapped for a 900 m range in both landscapes: from 3,500 m to 4,400 m in Huancavelica and 3,200 m to 4,100 m in Pasco. Across cultivar groups and landscapes, bred varieties occupied the widest altitudinal distribution of 1,100 m while bitter landraces had a narrow range of 400 m in Pasco. Bitter landraces began to occur at 3,800 m vs. 3,600 m of altitude in Huancavelica and Pasco respectively. All three cultivar groups overlapped between 3,800 m and 4,400 m in Huancavelica and 3,600 m and 4,000 m in Pasco.

**Fig 4.**
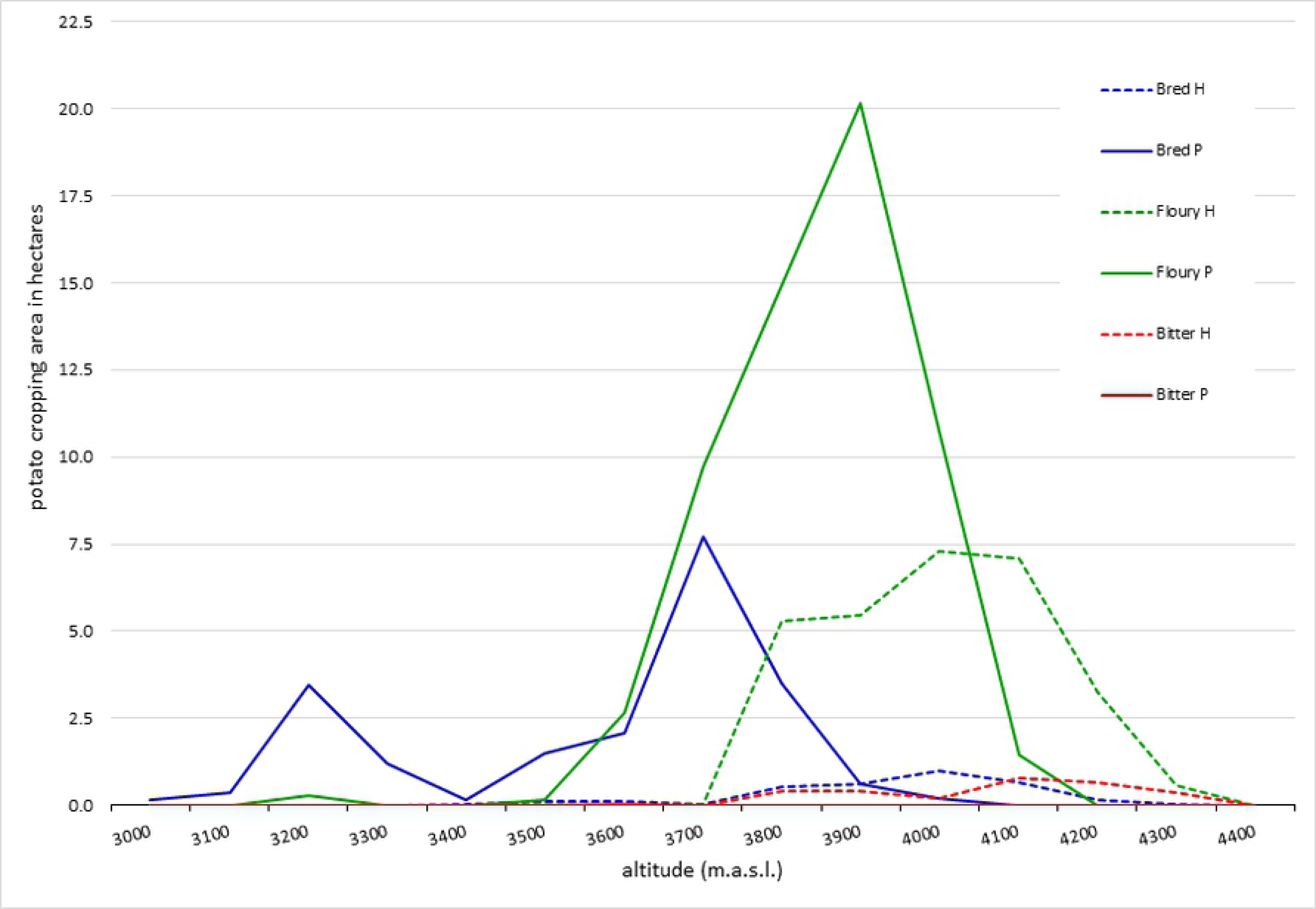
Total potato cropping area by cultivar group (bred, floury, bitter) and landscape (Huancavelica = H, Pasco = P) across the altitudinal range from 3000 to 4400 m.a.s.l.

We also examined the number of cultivars per field for incremental 100-meter altitudinal belts in each landscape. In Huancavelica, the highest concentration of cultivars occurred at the 4,000–4,100 m altitudinal belt with an average 37.0 (±12.7) and maximum 46 cultivars per field. These were floury, bitter and bred cultivars. This was the case at 3,900–4,000 m with an average 22.3 (±11.6) and maximum 50 cultivars per field in Pasco, involving only floury landraces and bred varieties. The highest levels of within-field diversity are concentrated at the upper limits.

### 3.5. Temporal characteristics of intraspecific diversity

#### 3.5.1. Fallow in rotations

Of 1,101 surveyed fields, 92.4% had a fallow period in the rotation. Remaining fields were cultivated uninterruptedly. The average period was a total of 7.4 years, either continuous or with one year of potato cultivation between two resting periods, for the ten-year cropping cycle recalled in the study. Fields with a fallow in the rotation represented 96.3% of fields in Huancavelica and 89.4% in Pasco. Average field-level fallowing rates were calculated for exclusive and mixed fields by cultivar group (Table 10). Fields containing exclusively bred varieties in Pasco showed the lowest fallowing rates (4.4 out of 10 years) and most intensive management compared to fields exclusively containing floury landraces (8.3 out of 10 years). Therefore, discriminatory management for fields with exclusively bred varieties or landraces occurred in Pasco. This was not the case in Huancavelica, where differences in fallowing periods between cultivar groups were smaller: 7.5, 7.4 and 7.2 years for fields containing bred varieties, floury and bitter landraces respectively. In both landscapes, we found a significant positive relationship (p<0.001) between the fallowing rate and altitude of fields (Fig 5A, 5B). The duration of fallowing periods tended to increase with altitude. However, in Pasco this relationship was stronger (R=0.35) compared to Huancavelica (R=0.12).

**Table 10.** Average fallowing rates for exclusive and mixed fields by cultivar group and landscape.

| Cultivar group | N† | Huancavelica |  |  |  |  | Pasco |  |  |  |  |
| --- | --- | --- | --- | --- | --- | --- | --- | --- | --- | --- | --- |
|  |  | Av. | Max. | Min. | SD. |  | N† | Av. | Max. | Min. | SD. |
| Bred varieties | 33 | 0.75 | 0.90 | 0.33 | 0.17 |  | 168 | 0.44 | 0.90 | 0.00 | 0.37 |
| Floury landraces | 235 | 0.74 | 0.90 | 0.00 | 0.19 |  | 376 | 0.83 | 0.90 | 0.50 | 0.07 |
| Bitter landraces | 21 | 0.72 | 0.90 | 0.50 | 0.10 |  | - | - | - | - | - |
| Mixed (BR+FL)‡ | 111 | 0.76 | 0.90 | 0.00 | 0.13 |  | 71 | 0.78 | 0.90 | 0.00 | 0.19 |
| Mixed (FL+BL)‡ | 55 | 0.69 | 0.90 | 0.00 | 0.23 |  | 4 | 0.89 | 0.90 | 0.88 | 0.01 |
| Mixed (BR+FL+BL)‡ | 26 | 0.67 | 0.90 | 0.00 | 0.26 |  | 1 | 0.80 | 0.80 | 0.80 | - |

†Number of fields for each exclusive and mixed cultivar group type.
‡BR=bred varieties, FL=floury landraces, BL=bitter landraces.

**Fig 5.**
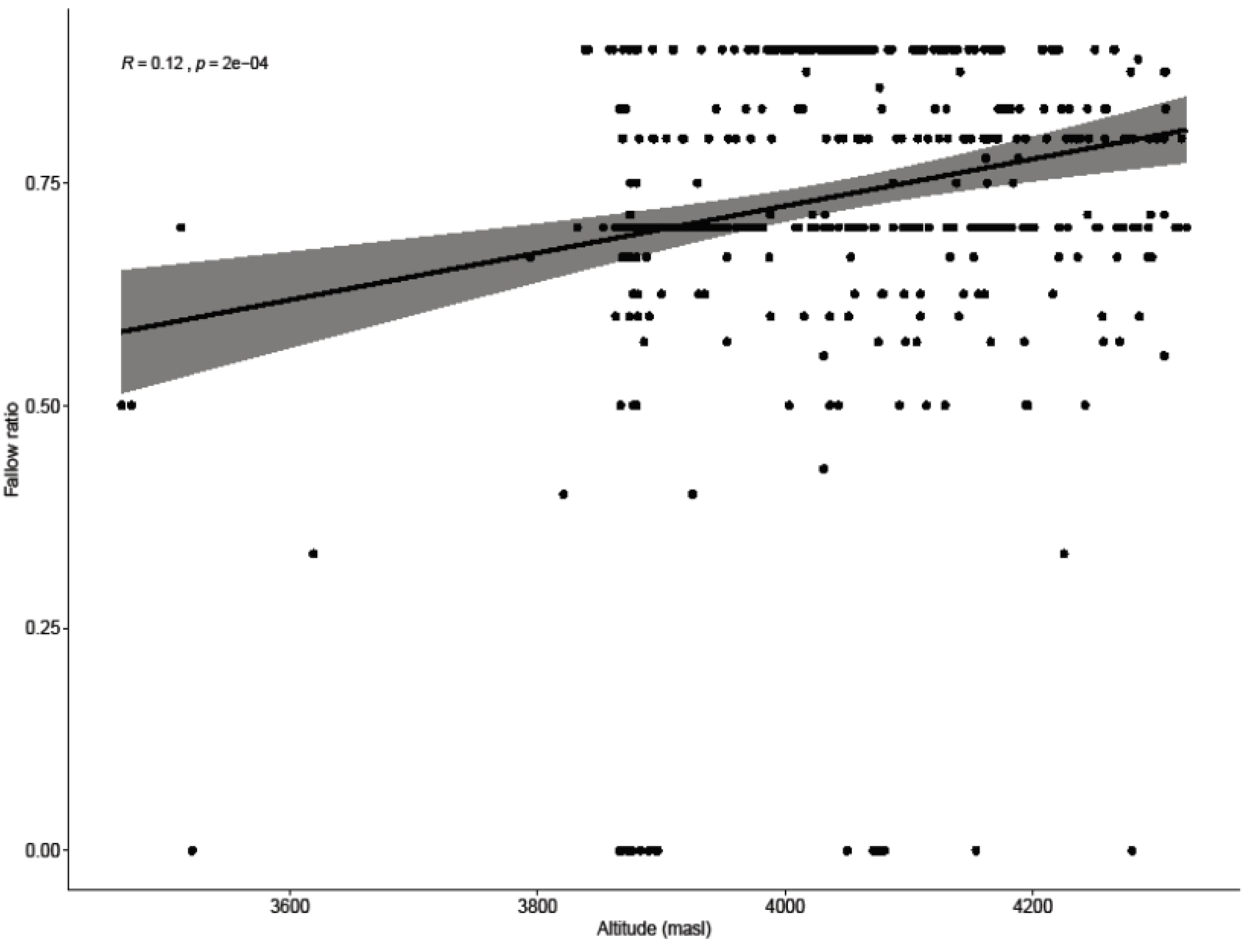
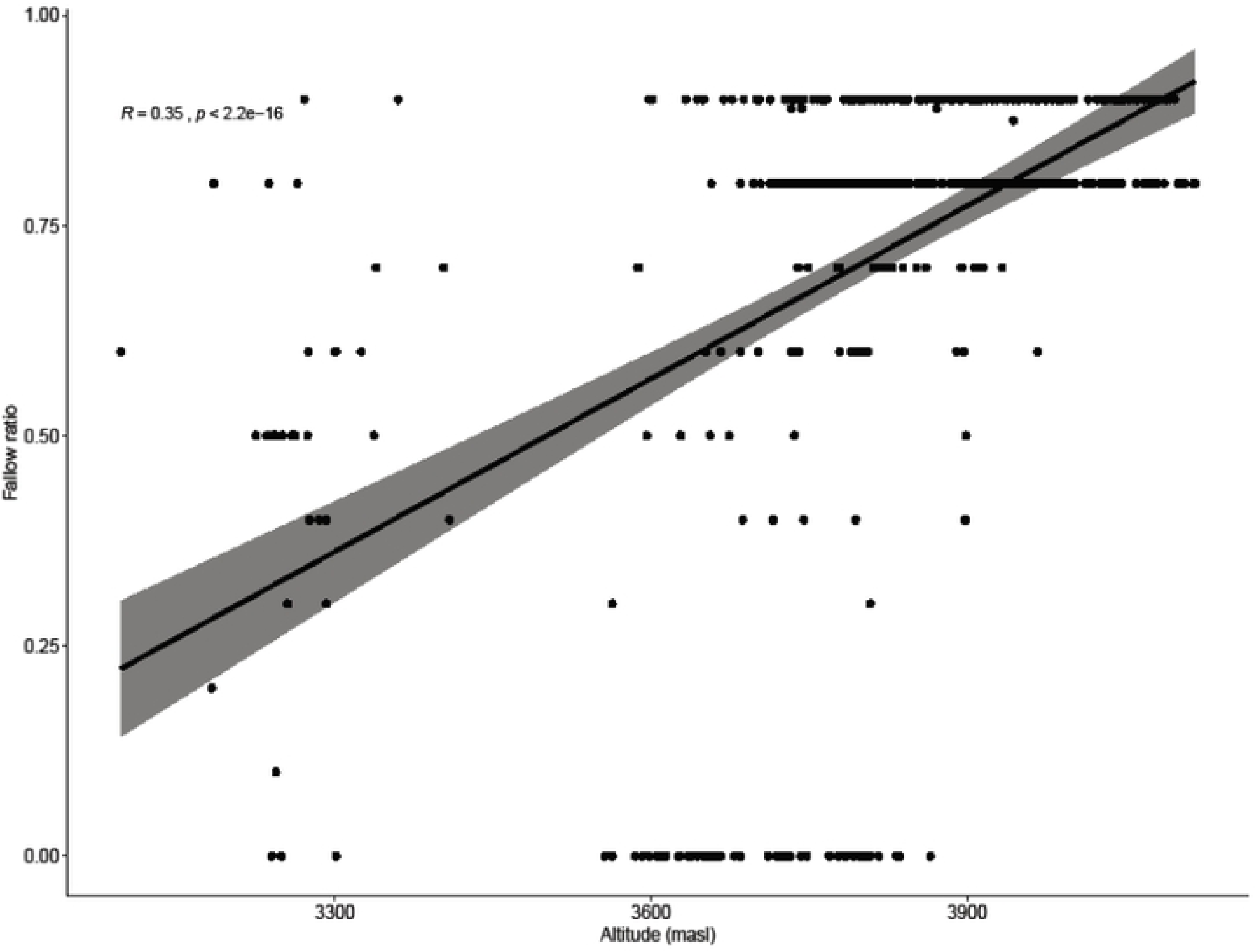
Significance of relationship between fallowing rate and altitude in the Huancavelica (A) and Pasco (B) landscapes.

#### 3.5.2. Rotation sequences

Most fields involved only potato in their cropping sequences: 54.1% in Huancavelica and 98.9% in Pasco. In Huancavelica, 7.3% of these fields involved two cultivar groups into their rotations, i.e. a bred varieties–floury landraces or floury landraces–bitter landraces sequence, and subsequently a fallow period. Remaining fields exclusively involving potato in this landscape obeyed the sequence bred varieties–fallow (6.5%), floury landraces–fallow (51.2%), bitter landraces-fallow (2.3%) and 32.7% involved mixed cultivar groups followed by a fallowing period. In Pasco, 10.3% of fields exclusively involving potato did not include a fallowing period in the cropping rotation. These were either uninterrupted bred varieties– floury landraces sequences (8.5%) or entirely dominated by bred varieties (1.8%). In this landscape, 16.1% of fields exclusively involving potato included bred varieties and floury landraces as mixed plots in a cropping sequence with a fallow, while 13.1% and 60.5% had a bred varieties–fallow and floury landraces–fallow sequence respectively.

Rotation sequences with other crop species were more varied and frequent in Huancavelica than Pasco at both intermediate and high altitudinal ranges (Fig 6). In Huancavelica, 44.5% of potato fields integrated cereals (oats, barley), 1.2% legumes (faba, lupine), 1.2% grasses (*Lolium multiflorum*), and 0.6% minor Andean tubers (*Ullucus tuberosus*, *Tropaeolum tuberosum*) in the rotation. Cereals were not included at all in rotation sequences with the potato in Pasco, and only 1.0% of fields incorporated a legume (peas) and 0.2% an Andean tuber (*Tropaeolum tuberosum*). Cereals were planted after floury landraces (20.8%), bitter landraces (2.7%), bred varieties (2.5%) and fields containing mixed cultivar groups (18.3%) in Huancavelica. Legumes in this landscape were planted after floury landraces (0.2%), bred varieties (0.6%) and mixed bred and floury cultivars (0.4%). All cropping sequences containing legumes and Andean tubers in Pasco occurred after bred varieties.

**Fig 6.**
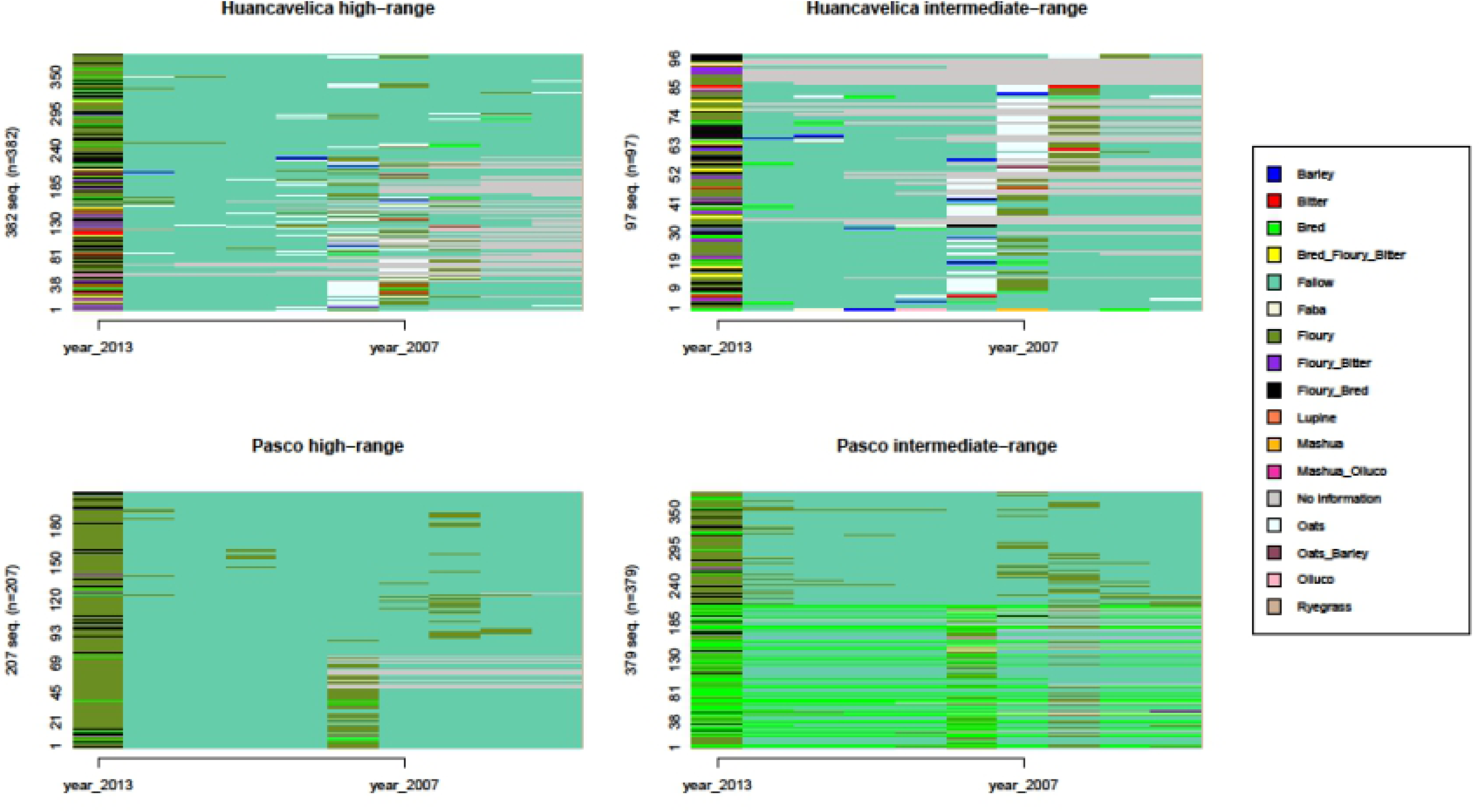
Pattern analysis of cropping sequences for intermediate and high-range altitude fields in each landscape.

#### 3.5.3. Association of fields with sectoral fallowing systems

Fields associated with a communal sectoral fallowing system comprised 32.4% of all surveyed fields and 33.5% of the total potato cropping area in Huancavelica. In Pasco, they represented 89.2% of fields and 92.1% of its total potato cropping area. The total area with potato under sectoral fallowing was 11.7 ha in Huancavelica and 74.5 ha in Pasco. These were covered 84.7% with floury landraces, 7.1% with bred varieties, and 8.2% with bitter landraces in Huancavelica. The potato cropping area under sectoral fallowing in Pasco was 80.5% floury landraces, 19.5% bred varieties and 0.04% bitter landraces. Areas that were not part of a sectoral fallowing regime comprised 23.3 ha in Huancavelica and 6.5 ha in Pasco. These were allocated 82.0% floury landraces, 10.2% bred varieties and 7.8% bitter landraces in Huancavelica; and 1.6% floury landraces, 98.4% bred varieties and 0.0% bitter landraces in Pasco. One hundred (100) of 130 cultivars in Huancavelica and 189 of 191 cultivars in Pasco occurred in areas under sectoral fallowing. Areas that were not managed as part of a sectoral fallow contained 105 cultivars in Huancavelica and 25 in Pasco.

In each landscape, we compared fields associated and not associated with sectoral fallowing systems for cultivar diversity per field, field size, and altitude. We identified significant and opposing differences in the altitudinal distribution of fields associated and not associated with sectoral fallowing systems. While in Huancavelica fields in sectoral fallows had a significantly lower median value in altitude compared to those outside such sectors (3,938 (±94) m vs 4,090 (±134) m, W=8823, p=2.2e-16), in Pasco, fields in sectoral fallows had a significantly higher median altitudinal value than fields dissociated from sectors (3,836 (±175) m vs. 3,679 (±145) m, W=30302, p=2.2e-16). No significant differences (p>0.05) were observed in cultivar diversity and field size between fields associated and not associated with sectoral fallows in Huancavelica. However, significant differences were observed for the same in Pasco. Sector fields had higher median values with respect to the total number of cultivars (5.9 (±7.6) vs. 1.4 (±2.5) cultivars per field, W=27582, p=4.481e-12) and field size (1,348 (±1,555) m² vs. 958 (±1,235) m², W=23107, p=0.0009386) in comparison to non-sector fields.

The sectoral fallowing sectors in Pasco were specifically targeted to landraces concentrating high levels of cultivar diversity while the non-sectoral fallowing land, subject to household-level decision-making, was predominantly destined to bred varieties and a limited number of commercial landraces in comparatively smaller field areas. Such a pattern does not show for Huancavelica where areal arrangements for cultivar group portfolios and cultivar diversity are evenly distributed across the two land-use systems.

### 3.6. Landscape differences by ‘fixed’ altitudinal ranges

Based on the generalized linear model (with elastic-based penalization) (see Materials and methods, section 2.6), we identified characteristics that significantly differentiated the management of intermediate-range fields (3,500 m to 3,899 m) across Huancavelica and Pasco. Product end use, tillage type, and mixed-cultivar fields were the top differentiators for this altitudinal range (Fig 7A; S1 Fig A, B, C). Intermediate-range fields in Pasco were significantly associated with production for sale (65% of fields), while in Huancavelica it was consumption as end use (95% of fields). Further, intermediate-range fields in Huancavelica were significantly associated with mixed-cultivar groupings containing floury and bitter landraces (12 % of fields), in contrast to Pasco, where less than 0.1% of its fields at this range showed this cultivar combination. Tillage type also differentiated the landscapes significantly, with all fields in Pasco being managed through *chiwa* tillage. In Huancavelica, 82.5%, 10.3% and 7.2% of fields at this range were tilled using *chiwa*, *chacmeo* and *barbecho* respectively.

**Fig 7.**
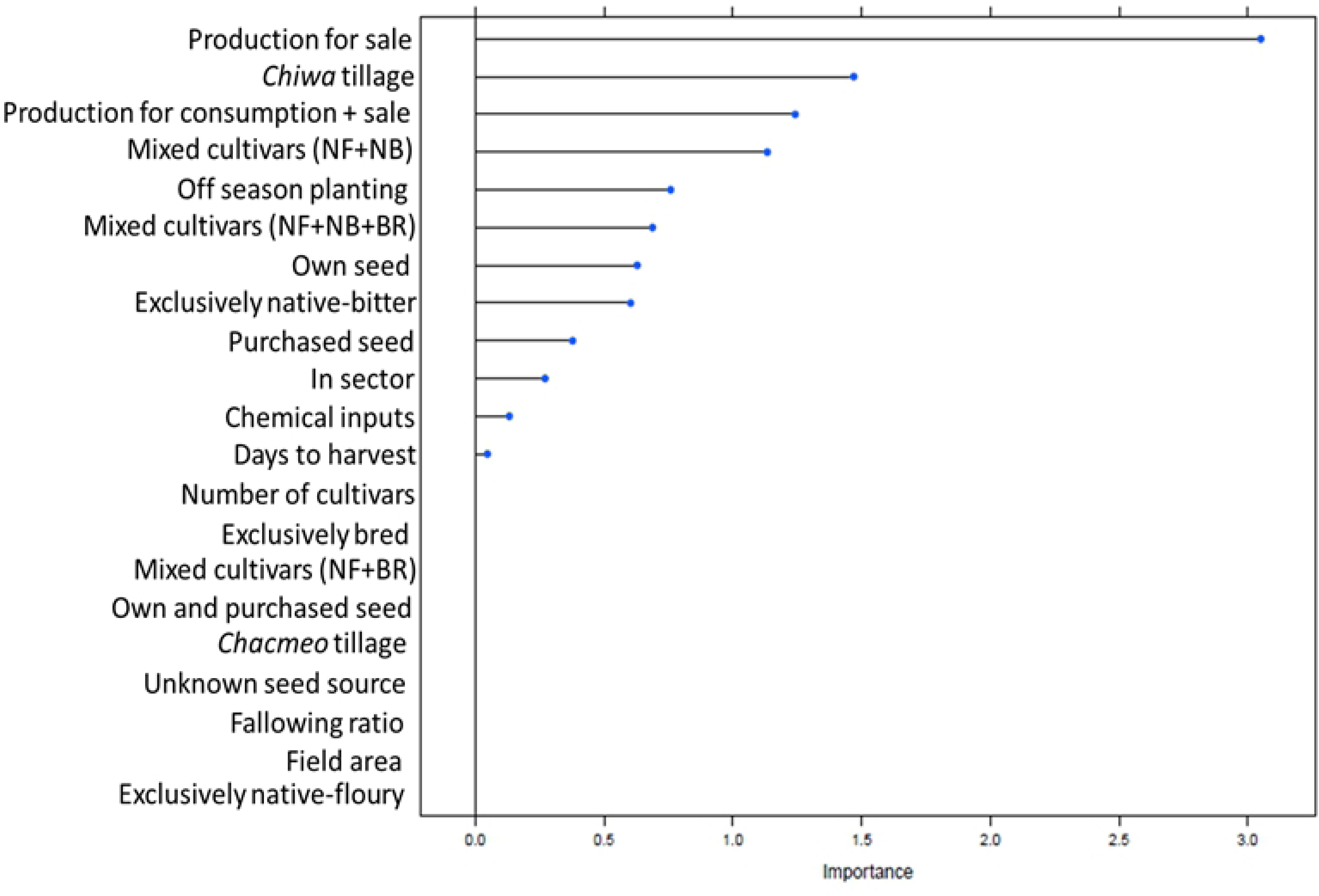
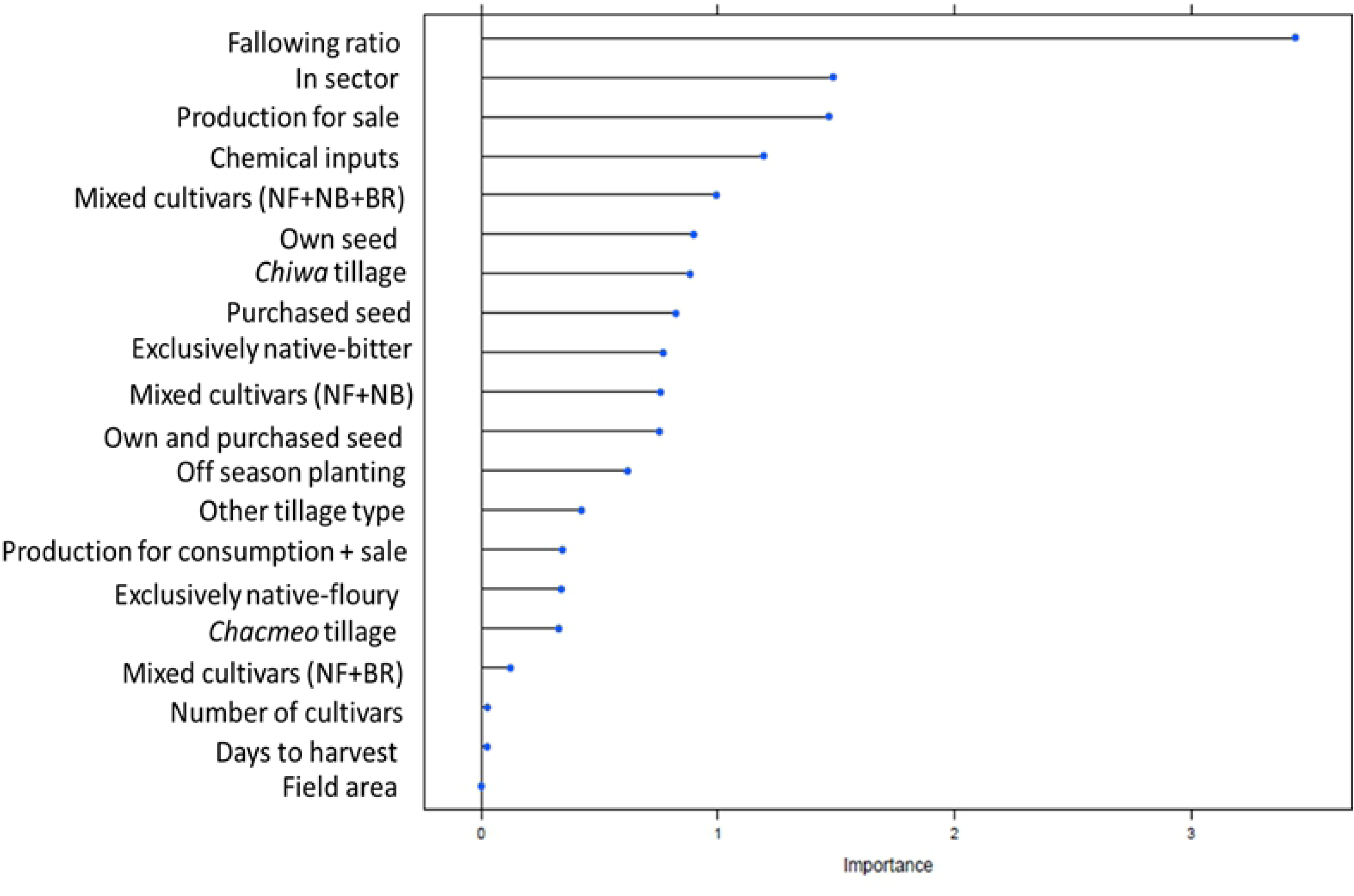
Farmer management associated variables listed in order of significance in differentiating (A) intermediate and (B) high-altitude fields between landscapes.

Analysis of upper-range fields (3,900 m to 4,324 m) revealed that fallowing ratio, number of fields associated with sectors, product end use, and chemical inputs were the top differentiating features of potato production between landscapes (Fig 7B; S1 Fig D, E, F). All fields in Pasco belonged to a fallowing sector. This applied to 23.3% of fields in Huancavelica. Field fallowing rates were also higher in Pasco at this range, 0.85 (±0.06) vs. 0.76 (±0.15) in Huancavelica. A significantly higher proportion of high-range fields (50%) was associated with sale in Pasco, in contrast to Huancavelica where significantly more fields (73%) were destined to consumption. Chemical inputs characterized all high-range fields in Pasco but only 31.9% of fields in Huancavelica. Seed source further significantly differentiated upper-range fields between landscapes, with farmers’ own seed applying to 99.7% of high-range fields in Huancavelica and 49.3% of fields in Pasco. In addition, high-range fields containing all cultivar groups occurred only in Huancavelica.

### 3.7. A timeline comparison of altitudinal distribution

The average altitudinal distribution of potato landraces in the two landscapes examined in this study has shifted upward by 330 m for floury landraces and 102 m for bitter landraces when comparing current ranges with those of passport data from the 1975–1985 genebank collection (Table 11; Fig 8; Fig 9). Pasco showed the greatest upward shift of 404 m for floury landraces. For bitter landraces, the upward shift has been less pronounced overall. However, in Huancavelica bitter landraces still showed a shift of 174 m. This contrasts with Pasco, where this cultivar group has, on average, moved upward by 31 m, although these results were obtained from a small number of samples.

**Table 11.** Altitude of landraces from 1975 to 2013 in the Huancavelica and Pasco landscapes.

| **1975-1985** | **N†** | **Av.** | **Max.** | **Min.** | **SD.** |
| --- | --- | --- | --- | --- | --- |
| Huancavelica | 31 | 3811 | 3973 | 3025 | 174 |
| Floury landraces | 29 | 3801 | 3973 | 3025 | 176 |
| Bitter landraces | 2 | 3948 | 3948 | 3948 | 0 |
| Pasco | 32 | 3519 | 3913 | 3135 | 165 |
| Floury landraces | 27 | 3494 | 3641 | 3135 | 156 |
| Bitter landraces | 5 | 3658 | 3913 | 3475 | 159 |
| **2012-2013** | **N** | **Av.** | **Max** | **Min** | **SD** |
| Huancavelica | 3323 | 4056 | 4324 | 3464 | 133 |
| Floury landraces | 2929 | 4057 | 4324 | 3521 | 128 |
| Bitter landraces | 153 | 4122 | 4324 | 3521 | 164 |
| Pasco | 3387 | 3883 | 4116 | 3097 | 125 |
| Floury landraces | 3132 | 3897 | 4116 | 3264 | 104 |
| Bitter landraces | 5 | 3829 | 3944 | 3646 | 117 |

†Number of reference cultivar samples.

**Fig 8.**
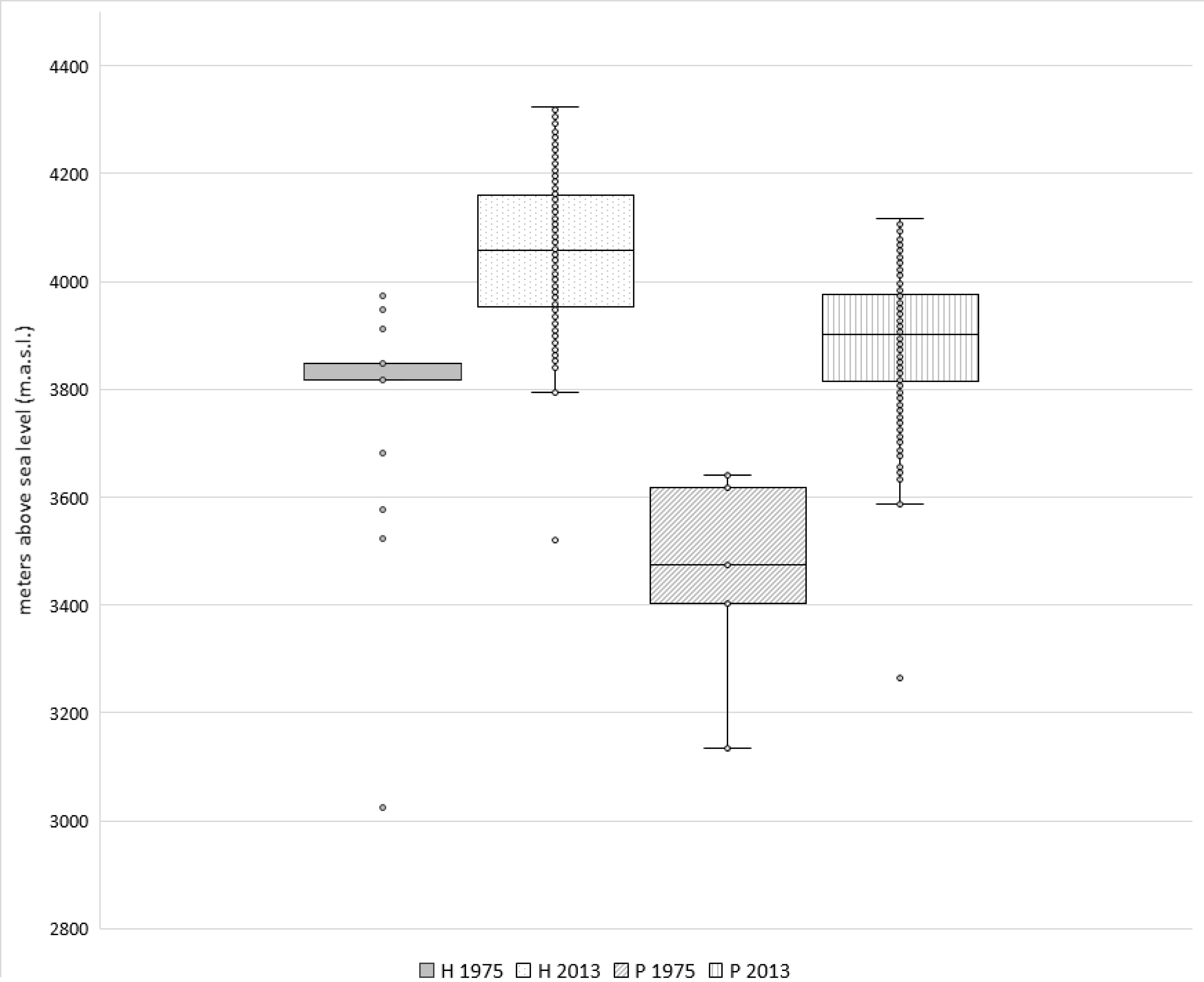
Altitudinal distribution of floury landraces (1975-2013) in m.a.s.l. (H = Huancavelica, P = Pasco).

**Fig 9.**
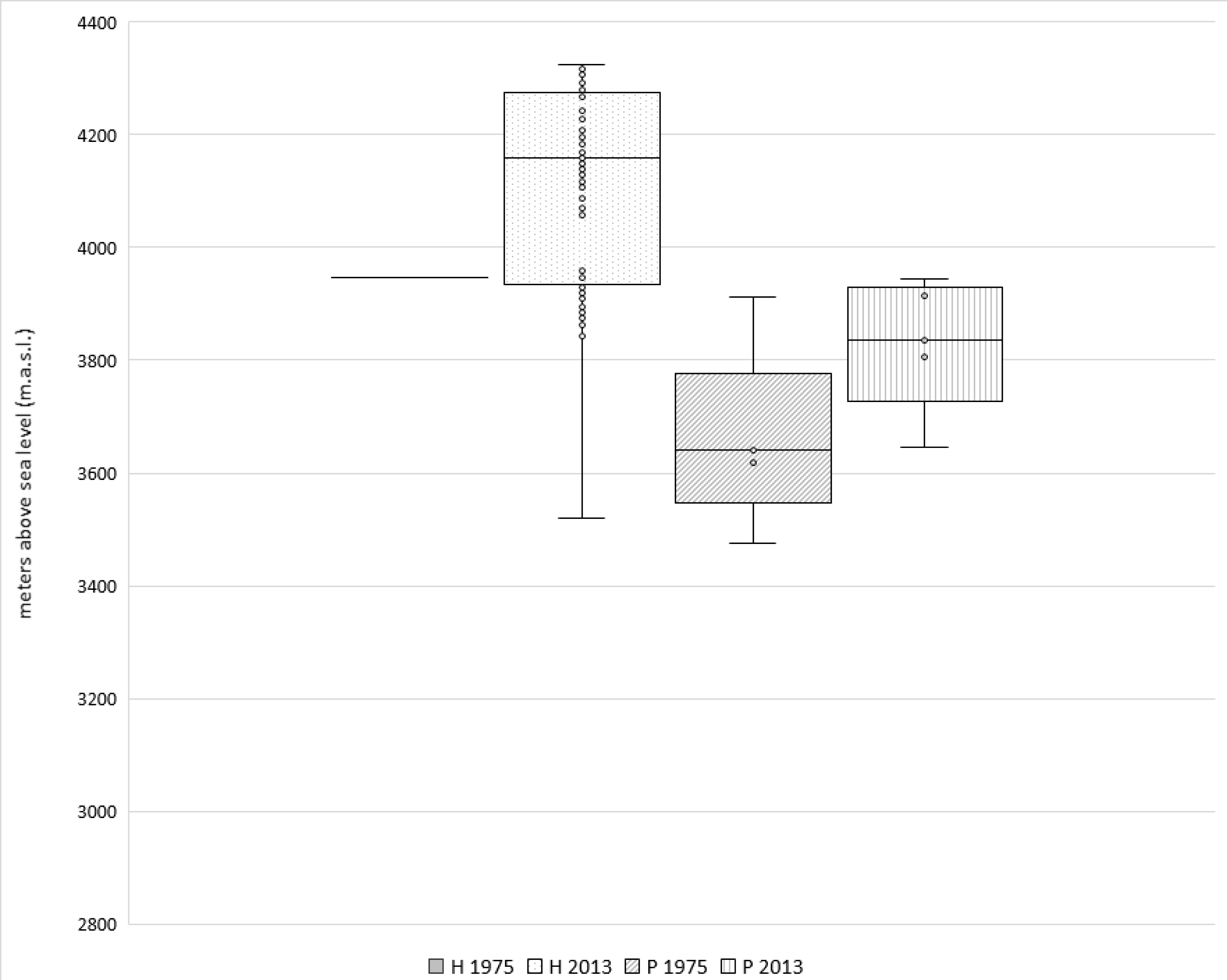
Altitudinal distribution of bitter landraces (1975-2013) in m.a.s.l. (H = Huancavelica, P = Pasco).

Maximum and minimum altitudinal distribution values also showed notable changes. The maximum reported altitude for floury landraces has increased by 475 m in Pasco and 351 m in Huancavelica. For bitter landraces in Huancavelica the shift in maximum altitude has been 376 m. As to minimum altitudes, floury landraces showed the highest increase by 496 m in Huancavelica. In Pasco the minimum altitude recorded for floury landraces has risen by 129m. The minimum altitude recorded for bitter landraces was surprisingly 427 m lower in 2013 than in 1975-1985 in Huancavelica, but it has shown a 171 m increase in Pasco.

## 4. Discussion

### 4.1. Hybrid landscapes and smallholder intensification

Our results show that smallholder land-use systems are spatially and temporally versatile, incorporating adaptations of traditional management practices to facilitate intensification. Such modifications of Andean cropping system components, allowing for the need to accommodate environmental and socio-economic pressures, have also been described by others [2, 21, 30, 31]. Intensification is occurring in its most basic form through shortening of fallow periods, but differently in each landscape. In Pasco, farmers ensure their ongoing production for both market and consumption by shortening the fallow period in their low-altitude fields while simultaneously maintaining long recovery periods in the upper-altitude range where most of the intraspecific diversity is also concentrated. The better household-level availability and access to land compared to Huancavelica enables farmers to manage their resources differentially and sustain commercial production of a few commercial cultivars while conserving diverse landrace portfolios at high altitude. In Huancavelica, on the other hand, the comparatively shorter fallow periods across all fields relate to diminishing land availability in a context of demographic pressure. With twice as many children and one third the total potato cropping area compared to Pasco, the only options that households have in this landscape involve shortened fallows and expanded cultivation at increasingly high altitudes [56, 67]. Adaptations become a necessity in contexts where land scarcity, the need for cash income from agriculture, and increased market orientation drive smallholder land use decisions [27, 68].

Hybrid land-use systems that integrate traditional and modern practices are common as smallholders adjust to changing production conditions and livelihood prospects in different ways [28, 69, 70]. This is notable in Pasco where, despite market-oriented intensification, two traditional land-use management components are more strongly maintained compared to the subsistence-oriented land-use systems of Huancavelica. Firstly, potato tillage in Pasco involved only the *chiwa* minimal-tillage system. This practice is common to sloping and high-altitude farming environments where the traditional foot plough or *chakitaklla* is typically used instead of animal or mechanical traction [56, 71]. A plausible explanation is erosion prevention on steep slopes under high rainfall conditions. Secondly, 92.1% of Pasco’s potato cropping area belonged to communal sectoral fallowing systems compared to only 33.5% of Huancavelica’s area. Intensification clearly hasn’t led to the disintegration of communal fallows.

Farmers in Pasco resorted to renting fields. This is only possible if land becomes available from households that have either migrated or oriented labor toward off-farm employment. Income generation through non-agricultural activities characterizes rural livelihoods across the Andes [1, 8, 11, 14]. Therefore, commercial agriculture partly drives intensification in Pasco. This is reflected not only in the low fallowing rates for fields where cultivation with bred varieties for sale is a priority but also by the consistent application of external inputs (fertilizers, fungicides) by all households. The use of chemicals can be partially attributed to high levels of late blight pressure. Except for a few bred varieties, most cultivars are actually highly susceptible to the disease [72, 73]. In contrast, in Huancavelica’s subsistence-oriented production systems, fallowing rates were particularly influenced by altitude, and the use of chemicals was very modest.

Huancavelica displays its own form of smallholder intensification in response to change. The traditional management of fields through communally coordinated sectors has to a large extent disintegrated and been replaced by cropping rotations that are directly decided upon at the household level. The disintegration and adaptations of sectoral fallowing systems have been documented throughout the Andes [30, 31, 45, 48, 74]. They are often a result of population growth, land scarcity, and the micro-fragmentation of landholdings, but have also been observed where access to irrigation provides smallholders with other crop production options [12, 67]. Soil degeneration and socio-cultural factors such as interrupted transmission of knowledge and discontinuity of communal decision-making institutions may also play a role [75, 76].

### 4.2. Conservation of landrace diversity amidst market specialization

A major driver of land-use change relates to economic integration and the consequent requirement for smallholders to specialize [77–79]. This tendency has previously been associated with diminished levels of crop varietal diversity [80–82]. In this study, we demonstrate that more subsistence-oriented agriculture does not necessarily encapsulate the highest landrace diversity. The commercial potato production in Pasco, which requires the adoption of intensive management practices, does not exclude parallel landrace conservation. These findings contrast with those reported in Ecuador by Skarbø (2014), who found a positive association between subsistence farming, Kichwa ethnicity and language, and the landrace richness of maize (*Zea mays*), common beans (*Phaseolus vulgaris*) and potatoes (*Solanum* spp.). Smallholders in Pasco, mostly mestizo Spanish speakers, are market-oriented producers of ware potato, particularly of bred varieties and commercial floury landraces. These smallholders intended the production of two-thirds of their total fields exclusively for sale, and consistently interacted with traders at the Carhuamayo market. In contrast, in Huancavelica only about one-fifth of fields were dual-purpose—destined to both consumption and sale—with the remainder being exclusively stored for home consumption. Yet, in Pasco, the total landrace diversity observed at the household and landscape levels was higher compared to Huancavelica. Market specialization and the allocation of significant areas to bred varieties does not displace landrace diversity, as Zimmerer (2013) also evidenced in Bolivia, where cash crop intensification and maize (*Zea mays*) agrobiodiversity were found to co-occur in smallholder landscapes.

Conversely, subsistence-oriented production accommodated more bred varieties in Huancavelica than in Pasco. Both as household-level average and as proportion of their collective cultivar diversity, more bred varieties were present in Huancavelica. Although not strictly market-oriented, smallholders in Huancavelica have integrated modern breeds into their portfolios due to their comparative advantage in terms of earlier maturation—which makes food available during the lean period—and ample accessibility in seed networks [57, 83]. This occurs even as the average cropping area per household is nearly three times smaller in Huancavelica than in Pasco. Here, predominantly indigenous Quechua-speaking smallholders don’t generate excess production for sale but maintain diversified cultivar portfolios with a higher representation of bred varieties and bitter landraces. In terms of areal coverage, there is more land available for diversity in Pasco. While proportionally Pasco’s diversity was grown on a smaller fraction of the household’s total potato area, in absolute terms the area occupied by landraces per household was nearly twice as large compared to Huancavelica. On the other hand, in Pasco more landraces were scarce or very scarce as they occupied a small proportion of the total cultivar portfolio. This can be partially explained by the way farmers allocate land and prioritize labor to generate an income. However, environmental factors likely also play a crucial role.

The source of seed tubers was almost entirely (99.6%) farm-saved in Huancavelica, but in Pasco this was only the case for 52.9% of fields. The extremely high altitudes at which potato cultivation occurs in Huancavelica are favorable for preventing virus infection and assuring seed health [84, 85]. Pasco, in contrast, is a high-risk zone for late blight disease and farmers mentioned seed quality as a continual concern. Seed degeneration resulting from cumulative pathogen and pest infestation over successive cropping cycles detrimentally affects yield performance and easily spreads across smallholder Andean networks [86]. Farmers in Pasco partially renew their seed stocks frequently by sourcing from higher-altitude production zones that meet their perceptions of quality for floury landrace production [57, 87]. With climate change, pest and disease pressure is likely to increase, warranting continuous monitoring of seed security and the conservation status of landrace diversity in both landscapes.

### 4.3. Uneven contemporary spatial distribution of landrace diversity

Our findings show that high intraspecific diversity persists in each landscape and collectively in Peru’s central Andes, especially of floury landraces. Yet this diversity is unequally distributed across landscapes. It is mostly concentrated at extremely high altitudes between 3,900 m and 4,200 m above sea level. The field scattering, overlap between cultivar groups, and use of mixed portfolios between and within fields show remarkable environmental plasticity and organizational ingenuity. It involves a continued use of diversity to adapt to an unpredictable environment and multiple production objectives [39, 54, 88]. Nonetheless, farmers commonly only prioritize five to seven landraces to meet mostly consumption or market needs. Bred varieties, which are a minor portion of the total varietal diversity (6.1%), cover the widest altitudinal distribution range while most landrace diversity is concentrated in a very narrow altitudinal range. This finding, confirming earlier reports of this kind of altitudinal concentration [30], suggests that diversity is potentially vulnerable with pests and diseases ‘pushing’ landraces upwards to limits where abiotic stress is highest (frost, hail) and land use for cropping competes with livestock.

Bitter landraces, which are characterized by relatively low diversity, were assigned only minimal area and were generally absent from farmers’ fields. Their apparent disappearance from the portfolios of most farmers may be the result of decreasing labor availability (needed to process them into *chuño*), changing consumer behavior, and less predictable frosts (in June) [89, 90]. Clearly, bitter landraces are at risk of being lost. The conservation dynamics of this special cultivar group warrants closer attention as their genetic potential is key to future breeding strategies to cope with abiotic stressors [40]. Traditional fallowing systems or *laymis* have been reservoirs of high intraspecific diversity in the central Andes. Yet, landrace diversity is not restricted to fields in fallowing sectors. In Huancavelica, the landrace diversity is currently contained in a landscape matrix of fields under a non-traditional household-level rotation with low-input management. In Pasco, the bulk of farmers’ diversity continues to occur in communally coordinated sectoral fallowing system with discriminatory, intensive management driven by market integration and late blight disease pressure. The above shows that diversity is being maintained as part of dynamic and adaptive management strategies.

Across landscapes, cultivar groups were not spatially separated but rather overlapped and to a large extent shared the same space. This finding confirms that rationales other than niche adaptation drive farmers’ spatial management of intraspecific diversity [2, 88, 91]. Potato cultivation in the two landscapes studied has moved upward by an average of 306 m since 1975. The altitudinal shift is most dramatic for floury landraces. For this cultivar group, contemporary maximum and minimum altitudes are 475 m and 500 m above that reported 38 years ago according to CIP passport data from collections. The incursion of the potato into higher altitudes has been previously documented and is explained by the compounding effect of environmental and social factors [22, 29, 56]. Changes in temperature and precipitation patterns, and lower number of and more erratic frosts are affecting agriculture in the central Andes [92–94]. Higher incidence of pests and disease is associated with climatic variability and further driving crop cultivation into higher altitudes [3, 6, 95]. Soil degradation also increasingly affects productivity in smallholder contexts, where population growth is pushing land-use systems beyond their capacity and into the upper limits of where agriculture is possible [20, 75]. Potatoes and their upward movement represent the highest cropping globally. Their changing land-use dynamics requires closer attention to understand the trade-offs and limitations of further altitudinal range expansion.

### 4.4. Study limitations

Assessments of land-use change and agrobiodiversity ideally require systematic comparisons over long periods. Data availability for timeline comparison is a constant limitation. In this study, we used a detailed inventory based on participatory GIS to examine the current situation. Yet, it represents only one season and does not account for inter-seasonal variation. We recorded the application of chemicals per field (yes/no) but did not measure the frequency or amounts of fertilizers and fungicides used. We therefore have no way of providing a fine-grained comparison of this type of intensification within and across landscapes. Further, we used folk taxonomy and focus group meetings to derive a master list of unique cultivars within and across landscapes. This is an adequate but imperfect way of classifying diversity, since it does not attain the precision provided by morphological and molecular characterization. Lastly, the genebank passport data from 1975–1985 only allowed for comparisons of altitudinal ranges for a limited number of floury and bitter landraces, excluding bred varieties.

## 5. Conclusions

The land-use dynamics of potato agrobiodiversity in the highlands of central Peru demonstrates remarkable adaptability in response to modern-day pressures. This is based on smallholder modification of traditional practices. High intraspecific diversity is maintained in these mixed, hybrid land-use systems. In each of the landscapes, intensification is taking place in different and rather unexpected ways. Whether predominantly market or subsistence-oriented, smallholder households inform their land-use decisions by drawing from the changing dynamics of their agroecological and socioeconomic contexts, increasingly geared toward intensification, i.e. shorter fallowing periods and chemical applications. Importantly, land availability gives smallholder households a comparative advantage by simultaneously enabling potato landrace conservation and market production. When it comes to on-farm agrobiodiversity, attributing the onus of its persistence on smallholders’ fields to market specialization may obscure the role of the other demographic, social, and environmental factors inherent in global change. Driven by population growth and pest and disease pressure, potato cultivation has moved into the upper limits of where agriculture is possible as shown by the comparison of contemporary altitudinal distributions with those of CIP’s genebank collections nearly four decades ago. Its landrace diversity is now concentrated in a narrow, upward moving altitudinal belt. The plasticity shown by the potato and the adaptability of smallholder land-use systems do not necessarily confer them resilience into the future. To gauge the on-farm dynamics of the potato in its center of crop origin systematic and long-term monitoring will be crucial. Its *in situ* conservation warrants the exploration of other options, such as the creation of incentives for smallholders’ diversity to be valued and utilized by society at large. From this standpoint, the active involvement of urban consumers and new institutional stakeholders may be key to the ongoing use and conservation of the potato’s intraspecific diversity.

## Acknowledgments

The authors thank the smallholder farmers and community authorities of Castillapata, Huachhua, Pumaranra, Bellavista and Chupaca for collaborating with us and making this research possible. We are grateful to the local teams of field enumerators who supported us during the arduous data collection. We also thank the International Potato Center for facilitating the genebank collection data that enabled our comparative timeline series analysis.

**S1. Fig. Independence analysis (based on chi square statistical testing of Pearson residuals) of the top most differentiating variables between intermediate and high-altitude fields in the Huancavelica and Pasco landscapes.** A) Production end use, B) Tillage type, C) Cultivar combination (NF=Native-floury; NB=Native-bitter; BR=Bred; Mixed=combinations of NF, NB and BR) show that intermediate-range fields in Huancavelica were associated with production for consumption, *chacmeo* and *barbecho* tillage, and mixed-cultivar groups of floury and bitter landraces compared to fields in Pasco; D) Fallowing sector association, E) Production end use, F) and Fallowing rates show that high-range fields in Huancavelica were not associated with a fallowing sector, production end use was destined to consumption, and fallowing rates were significantly lower compared to their homologues in Pasco. The scale corresponds to Pearson residuals and the color on the scale corresponds to a significantly positive (blue) or significantly negative (red) relationship based on independence analysis at p-value < 0.05.

